# Reduced Dimension, Biophysical Neuron Models Constructed From Observed Data

**DOI:** 10.1101/2021.12.03.471194

**Authors:** Randall Clark, Lawson Fuller, Jason A. Platt, Henry D. I. Abarbanel

## Abstract

Using methods from nonlinear dynamics and interpolation techniques from applied mathematics, we show how to use data alone to construct discrete time dynamical rules that forecast observed neuron properties. These data may come from from simulations of a Hodgkin-Huxley (HH) neuron model or from laboratory current clamp experiments. In each case the reduced dimension data driven forecasting (DDF) models are shown to predict accurately for times after the training period.

When the available observations for neuron preparations are, for example, membrane voltage V(t) only, we use the technique of time delay embedding from nonlinear dynamics to generate an appropriate space in which the full dynamics can be realized.

The DDF constructions are reduced dimension models relative to HH models as they are built on and forecast only observables such as V(t). They do not require detailed specification of ion channels, their gating variables, and the many parameters that accompany an HH model for laboratory measurements, yet all of this important information is encoded in the DDF model.

As the DDF models use only voltage data and forecast only voltage data, they can be used in building networks with biophysical connections. Both gap junction connections and ligand gated synaptic connections among neurons involve presynaptic voltages and induce postsynaptic voltage response. Biophysically based DDF neuron models can replace other reduced dimension neuron models, say of the integrate-and-fire type, in developing and analyzing large networks of neurons.

When one does have detailed HH model neurons for network components, a reduced dimension DDF realization of the HH voltage dynamics may be used in network computations to achieve computational efficiency and the exploration of larger biological networks.

## 1 Introduction

### 1.1 General Setting

When we have observed data generated by a complex, nonlinear system, neurons and their networks are a prime example of this, but we do not have a model for the detailed neurodynamics of the system, it is possible to use those measured data alone to create a dynamical rule that forecasts the future of the observed quantities beyond the window in time where the data was measured. We call this Data Driven Forecasting (DDF).

It is our goal in this paper to demonstrate how to achieve, then utilize DDF in the context of neuroscience.

DDF is to be viewed as a way to capture observed aspects of neurobiological data in contrast to the usual method of creating a detailed biophysical model, often of Hodgkin-Huxley (HH) type, which may or may not be correct, specifying all of the relevant ion currents, the required gating variables, and the concentrations of relevant quantities such as [*Ca*^2+^](*t*) [18, 34].

Unknown quantities in such a HH model may be estimated using data assimilation (DA) [38, 21, 25, 27, 2]. DA requires a model of the observed complex system, and, of course, data from observing that system. The data are used to train items in the model such as fixed parameters and unobserved state variables.

DDF does not require a model of all these, often unobservable, details. Nonetheless, as it is built on observed data, it encodes those details while forecasting only the observable properties of the neuron activity. Both DA and DDF may be seen as a form of supervised learning [3]. In this regard they may also be regarded as methods of machine learning.

If the construction and analysis of a biophysically detailed HH model has been achieved, perhaps employing DA, using HH models in large networks of biological interest may prove computationally quite challenging. DDF may be utilized to accurately forecast the voltage time course of this HH model thus replacing it in the network of interest. Only the voltages across neuron cell membranes are used in the communication among neurons in a network, thus DDF neurons are nicely suited for use as the dynamical elements of functional biological networks. Using a reduced dimensional model in network studies can result in significantly decreased computational tasks.

### 1.2 Data of Interest

We start with the formulation of DDF models for individual neurons.

We have in mind data where a neuron is stimulated by a known forcing via an injected current *I_stim_*(*t*), and its membrane voltage response V(t) is measured; namely, current clamp experiments. One observes V(t) at discrete times *t_n_* = *t*_0_ + *nh*; *n* = 0, 1, …, *N*. We use these data to build a biophysically based nonlinear discrete time map that takes *V*(*t_n_*) → *V*(*t*_*n*+1_) for any selected stimulating current. Importantly, the map must predict well for *n* ≥ *N* + 1.

We demonstrate that this is accomplished in in the analysis of numerically generated data from a standard neuron model and in the analysis of current clamp data collected in a laboratory environment.

When this is successful, we will have created a DDF dynamical rule moving V(t) forward in time without regard for gating variable time courses, parameters such as maximal conductances or reversal potentials, chemical reaction rates, or any of the other detailed biophysical characteristics of the HH neuron dynamics. Yet, built from observed data, the biophysical information is embedded in the DDF model.

As we shall demonstrate here, one is able to accomplish this but, not surprisingly, must give up some things that are found in the use of a detailed model. The method is restricted to forecasting only what is observed.

### 1.3 Useful Attributes of DDF Neurons

This forecasting construction *V*(*t_n_*) → *V*(*t*_*n*+1_), which we call a ‘DDF neuron’, may be used in network models of interest. DDF neurons would be located at the nodes of the network leaving only the network connectivity to be determined, possibly by DA from observed network activity data [2].

In biological networks, the individual neurons are driven by external currents, if present, and by the currents received from other neurons presynaptic to it. The gap junction and synaptic current connections are described by the presynaptic voltage and the postsynaptic voltages allowing DDF neurons to be valuable, efficient network nodes in computational models of functional neural networks.

If one has the goal of controlling aspects of functional neural networks to achieve desired goals, DDF neurons provide a computationally inexpensive way of incorporating the observable properties of such a network. This is significant as observables are the attributes which control forces may affect.

## 2 Plan for this Paper

1. To begin we describe the DDF method as applied to individual neuron data and give an example using numerical data from solving a basic Hodgkin-Huxley (HH) neuron model [15, 18, 34] with Na, K, and Leak channels. For brevity we designate this HH model as an NaKL neuron.
2. The time courses for the voltage and the Na and K channel gating variables {*V*(*t*), *m*(*t*), *h*(*t*), *n*(*t*)} from the model are used as ‘data’ and analyzed in two settings:

- The first setting is a confidence building exercise, *not biologically realistic*, that assumes we have observed data on the membrane voltage V(t) as well as on all three of the HH gating variables {*m*(*t*), *h*(*t*), *n*(*t*)} in the model.
- The second setting conforms to what one can actually do in a current clamp experiment, namely observe only the membrane voltage V(t) given the stimulating current *I_stim_*(*t*). This requires us to add to the basic DDF formulation the idea of constructing enlarged state spaces from the observed variables and their time delays [37, 4, 5, 1, 20]. This method is familiar and essential in the study of nonlinear dynamics and will be explained in the present context.
3. We next turn to the DDF analysis of laboratory current clamp data acquired in the Margoliash laboratory at the University of Chicago.
4. An analysis is then made of how DDF neurons can be used in the construction and study of networks of neurons. In this paper we first direct our attention to a quite simple two neuron example with gap junction connections.
5. Then we present a study of a ‘network segment’ where a presynaptic neuron, driven by a stimulating current *I_stim_*(*t*), drives a postsynaptic neuron via a ligand gated synapse.
6. A Summary and Discussion completes the paper.
7. We include three Appendices:

- Appendix A is a brief manual discussing how one builds DDF models in practice.
- Appendix B has the formulation for incorporating ligand gated synaptic connections into a DDF based network model.
- Appendix C contains a short biophysical discussion on the choice of stimulating currents *I_stim_*(*t*) selected to permit the DDF model to generalize its response to a wide class of stimulating currents. This appendix was developed following questions from two reviewers of an earlier draft of this paper.

## 3 Discussion of the Methods of DDF

Our discussion begins with a broader formulation of DDF than will be required in neurobiology where only V(t) is observed in laboratory current clamp experiments. We return to the realistic scenario of observing only V(t) after providing the broad perspective. An example of data, beyond membrane voltage, where DDF will permit accurate forecasting includes fluorescence associated with Ca concentration variation [*Ca*^2+^](*t*) in the presence of a Ca indicator [33, 39, 23], [*Ca*^2+^](*t*). The DDF formulation for [*Ca*^2+^](*t*) is described in Section (7).

The idea of DDF is to start with the collection of D-dimensional observed data **u**(*t_n_*) = **u**(*n*) = {*u_a_*(*n*)}; *a* = 1, 2, …, *D*, sampled at discrete times *t_n_* over an observation window [*t*_0_ ≤ *t_n_* ≤ *t_N_*]; *t_n_* = *t*_0_ + *nh*; *n* = 0, 1, 2, …, *N* in time steps of size *h*.

Next, **using only these data**, we discuss how to construct a discrete time map **u**(*n* + 1) = **u**(*n*) + **f**(**u**(*n*), **χ**) for forecasting the future of those data. The **χ** are parameters in **f**(**u**(*n*), **χ**) that we will estimate (train) on the observed data. The nonlinear function **f** (**u**, **χ**) is called the *vector field* [36] of the discrete time map.

When we do not have a model of the biophysical processes generating **u**(*t*), we ask: can we build a *representation* of **f** (**u**, **χ**) and use observed data to determine the unknown, time independent quantities **χ** in that representation?

We will answer this in the affirmative using applicable developments in applied mathematics [13, 6, 8, 19, 32, 22] explored in depth over many years.

The general idea was well investigated in the context of autonomous systems where there is no external forcing. This is not the situation that one addresses in neurobiology where neurons are stimulated by external sources and, in functional networks, by the activity of other neurons in the network.

The first step is to select a parametrized representation for **f** (**u**, **χ**). Using **u**(*n* + 1) = **u**(*n*) + **f** (**u**(*n*), **χ**) for 0 ≤ *n* ≤ *N*, to estimate (train) the parameters **χ**. Once the **χ** are known, we are able to use this trained discrete time dynamical rule **u**(*n* + 1) = **u**(*n*) + **f** (**u**(*n*), **χ**) to forecast the behavior of the observed quantities **u**(*n*) for *n* ≥ *N* + 1 in time steps of size h.

We use the results of [32, 22] showing that one can accurately *represent a multivariate function of* **u**, **f** (**u**, **χ**), as

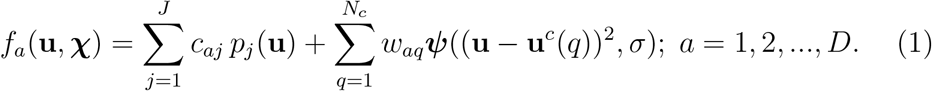

The {*c_aj_, w_aq_*, *σ*} are among the parameters to be estimated/trained using the observed data {**u** (*n*)}.

In Eq. (1) the *p_j_* (**u**) are multivariate polynomials of order j, The functions ***ψ***((**u** – **u**^*c*^(*q*))^2^, *σ*) are called *radial basis functions* (RBFs). The *N_c_* {*u^c^*(*q*)} are denoted as *centers*, and they are selected from the observed data; *N_c_* ≤ *N*.

In developing this representation of the vector field as a function on **u** space, we may think of the {**u**(*n*); *n* = 0, …, *N*} as samples of a distribution in **u**. The **f** (**u**, **χ**) are designed to interpolate among these samples

The training of the {*c_aj_*, *w_aq_*} is a linear algebra problem [29]. The linear algebra problem for *N_c_* ≠ *N* requires regularization, and that means we must specify a Tikhonov regularization parameter, which we call *β*. This is also called ridge regression.

We must also specify any parameters appearing in the functions ***ψ***((**u** – **u**^*c*^(*q*))^2^, *σ*). A guide to how we select all parameters of the DDF formulation, in practice, including {*c_aj_*, *w_aq_*, *σ*, *β*}, is given in Appendix A. When we use time delay coordinates, as we do in Section (6), two more parameters enter: the time delay τ and the dimension of the time delay vector *D_E_*. So **χ** = {*c_aj_, w_aq_*, *σ*, *β*, *τ*, *D_E_*} is the full set of parameters that we must estimate from the given data.

The representation of the vector field *f_a_*(**u**, **χ**) in Eq. (1) has both what one often finds described as RBFs by themselves (the second term on the right) and polynomials in **u**. We first tried to work with the polynomials alone, but found that when *J* was only 3 or 4, they were not able to represent the kind of nonlinearities found in biophysical models of neurons. As *J* increases the number of coefficients *c_aj_* grows more or less as *J*!, and even the linear algebra problem becomes difficult. The more general case in Eq. (1) is discussed in [32, 22]. In practice we only used the monomial **u**.

The possibility of including polynomial terms beyond **u**(*n*) was not required to achieve the results we present, so the training (estimation) of **χ** is done using

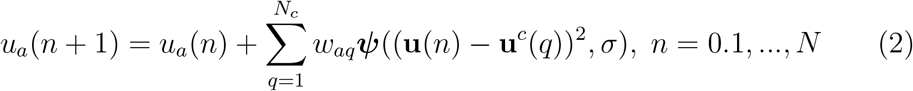

and we realize it by minimizing

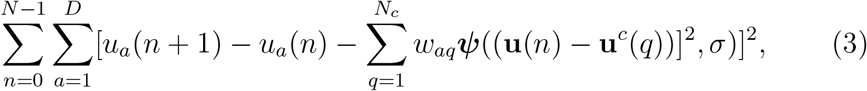

which we regularize in a well established way [29]. The details of this are presented in Appendix A.

Once the linear algebra problem of determining the {*w_aq_*} is completed Eq. (2) becomes our discrete time (in steps of size h) dynamical, nonlinear forecasting rule for times *t_n_* ≥ *t_N_*.

There are many choices for these RBFs [13, 22, 32, 7]. Our RBF choice has been the Gaussian: ****ψ***_G_*((**u** – **u**^*c*^(*q*))^2^) = exp[–*R*(**u** – **u**^*c*^(*q*))^2^)], *R* = 1/*σ*^2^. Other choices, and there are many, see Table I in [32], have given equivalent results in practice.

## 4 Using DDF in Neurobiology

### 4.1 Data Assimilation

In the study of the ingredients of functional neuronal networks, one is able to measure the time series of voltage V(t) across the cell membrane of individual neurons in a routine manner. Using observed values of V(t) along with knowledge of the forcing by a stimulating current *I_stim_*(*t*) presented to the neurons, it is often possible to estimate the detailed electrophysiological properties of a Hodgkin-Huxley model of an individual neuron [15, 34, 18] using data assimilation [2].

The data assimilation estimation involves inferring the unmeasured state variables, including ionic gating variables, and unknown parameters such as maximal conductances of ion channels. A model, developed and completed in this manner, is validated by comparing its voltage time course when driven by stimulating currents after the observation window. Only in the observation window is information passed to the model from measurements of V(t).

### 4.2 DDF

The DDF perspective selects a related, but also biophysically grounded, path that emphasizes what can actually be measured and forecasts only those aspects of complex neuronal activity.

Approaching the question of biophysical models for the dynamics of neurons from a DDF perspective provides a way to make predictions/forecasts of V(t) *without* the details of the biophysical model. This, as noted, sets aside the knowledge of the details of the model, but for purposes of building up the dynamics of voltage activity in a network, it may be of great utility.

One attractive feature of DDF is that it results in significant model reduction from the many state variables and proliferation of parameters present in the neuron dynamics [27] by focusing on those state variables that can be measured.

### 4.3 Hodgkin-Huxley Structure for Driven Neurobiological Dynamics

We continue our discussion of DDF in neurodynamics by attending to how one can work with numerical/simulated data from a basic, well studied, Hodgkin-Huxley (HH) model neuron. Our analysis of experimental current clamp data will follow this set of numerical examples.

The biophysical HH equations for the dynamics of a single neuron driven by a simulating current *I_stim_*(*t*) have the structure

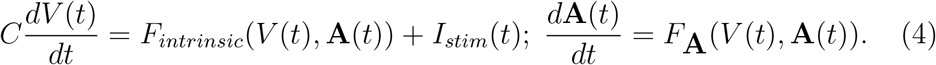

*F_intrinsic_* (**A**(*t*), *V*(*t*)) contains the ion currents and their gating variables which satisfy **A**(*t*), 0 ≤ **A**(*t*) ≤ 1. These quantities are descriptive of the intrinsic electrophysiology of a biophysical neuron are independent of the stimulating current *I_stim_*(*t*). In addition to the HH dynamics there may be other state variables in addition to {*V*(*t*), **A**(*t*)} such as concentrations of various biochemicals. To proceed, we concentrate on the HH voltage equation structure in Eq. (4).

In the HH formalism the equation for the voltage is driven by *I_stim_*(*t*) in an additive manner. We will use this in formulating our DDF protocol for biophysical neuron models. What we do not know from the data alone are the vector fields {*F_intrinsic_*(*V*(*t*), **A**(*t*)),*F*_**A**_(*V*(*t*), **A**(*t*))} whose specification yields the detailed HH biophysical model.

The solution or flow [36] of the HH model, Eq. (4), is achieved by integrating Eq. (4) from *t_n_* to *t_n_* + *h* = *t*_*n*+1_ leading us to the discrete time map 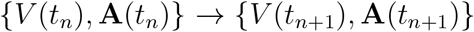 or 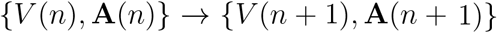:

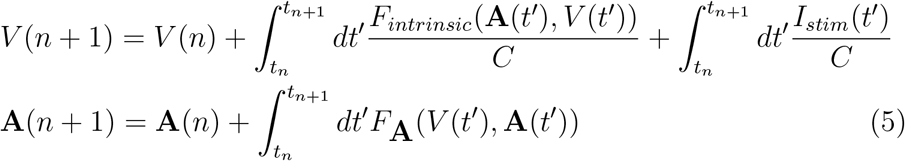

### 4.4 The Basic NaKL HH Neuron as An Example

We now analyze the DDF representation of the flow, Eq. (5) in the NaKL HH model neuron in two cases: (1) when we observe all state variables {*V*(*t*), **A**(*t*)}, and (2) when we observe only V(t). Case (1) is not a realistic scenario in current clamp experiments, but we include it as a confidence building exercise in the construction of DDF neurons. Case (2) is the realistic scenario in a current clamp experiment where a simulating current *I_stim_*(*t*) drives a neuron and only the membrane voltage V(t) is observed.

Our first step is to work with numerically generated data from the HH NaKL model neuron [18, 34].

This basic, well studied, detailed biophysical HH neuron model has 4 state variables and the order of 20 parameters [18, 38, 21]. DDF will replace this with a forecasting equation for V(t), the experimentally observable quantity, alone. As we have argued, much is gained by this model reduction.

The equations for this neuron model are

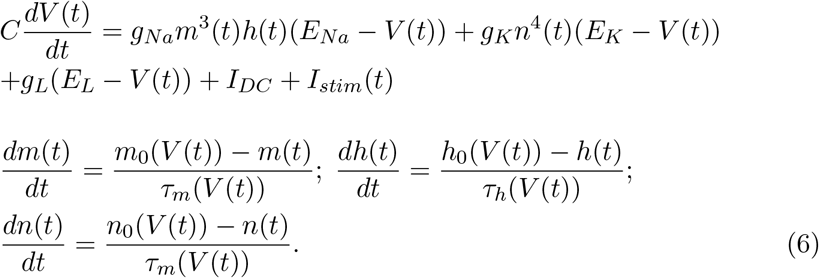

The gating variable functions *g*_0_(*V*), *τ_g_*(*V*); *g* = {*m*, *h*, *n*} have the form

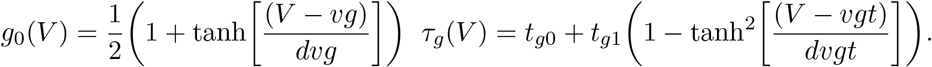

In these equations we use the parameters given in [38, 21]. Data is generated by solving Eq. (6) using a fourth order Runge-Kutta method [29].

The DDF formulation when we observe **w**(*t*) = {*V*(*t*), *m*(*t*), *h*(*t*), *n*(*t*)} = {*V*(*t*), **A**(*t*)}, *again* **not** *the realistic biological feature of a current clamp experiment*, is the following (**w**(*t_n_*) = **w**(*n*)):

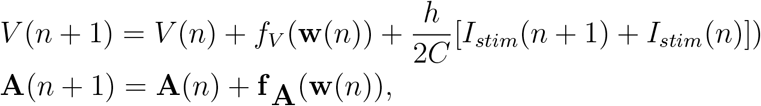

as suggested by Eq. (5). The functions {*f_V_*(**w**), **f_A_**(**w**)} are sums over the RBFs as in Eq. (1).

We use the trapezoidal rule [29] for the integration over *I_stim_*(*t*). We are taking steps of size h in time. There are many improvements over the trapezoidal rule for the integration from *t_n_* to *t_n_*+h over *I_stim_*(*t*). Those improvements require sampling or estimating values at points between *t_n_* and *t_n_* + *h*. We do not have these quantities in our data set.

The task for DDF is to select representations for the vector fields *f_V_* (**w**) and *f*_**A**_(**w**) which we shall do below. Interestingly, the constant C, the membrane capacitance, can be estimated in the DDF training protocol as the time course of the driving force, *I_stim_*(*t_n_*), must be specified by the user.

Using a Gaussian RBF for each of the four vector fields {*f_v_*(**w**), *f*_**A**_(**w**)}, DDF training will provide an estimate of the parameters in each vector field. The result from forecasting with the trained DDF model is shown in Fig. (1). It is clear from this result that the DDF when all {*V*(*t*), **A**(*t*)} are ‘observed’ does a strikingly good job at forecasting the actual observable V(t).

**Figure 1:**
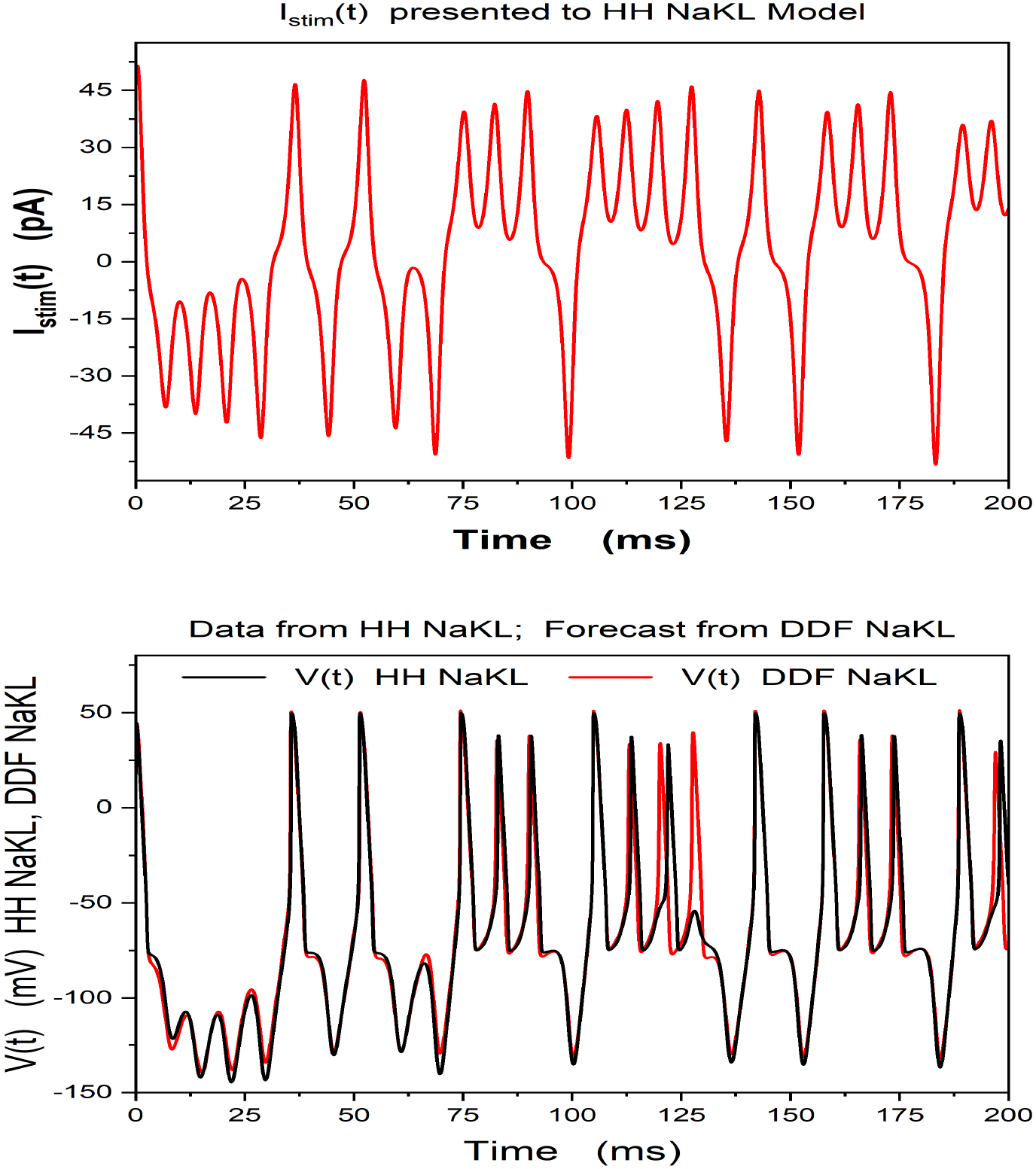
**Top Panel** Our selection of a stimulating current to an NaKL Hodgkin-Huxley neuron model [38, 21]. Data is generated using this *I_stim_*(*t*) in solving Eq. (6). **Bottom Panel** Forecast using a Gaussian RBF Model trained by **both the voltage and the gating variables:** {*V*(*t*), *m*(*t*), *h*(*t*), *n*(*t*)}. (`This not a realistic protocol for a current clamp experiment where` *I_stim_*(*t*) `is given, but V(t) alone is observed. This calculation is only a demonstration of the efficacy of DDF method in neurodynamics`.)

## 5 Observing only V(t): Time Delay Methods

What is actually measured in current clamp laboratory experiments is V(t) alone. The neuron dynamics resides in a higher dimensional space than the one dimensional V(t) that is measured. What we observe is the operation of the full dynamics projected down to the single dimension V(t). To proceed we must effectively ‘unproject’ the dynamics back to a ‘proxy space’, comprised of the voltage and its time delays [37, 4, 5, 1, 20], which is equivalent to the original state space of V(t) and the gating variables for the ion channels.

This is accomplished as follows: If we have observed V(t), we can define *D_E_*-dimensional (‘unprojected’) proxy space vectors **S**(*t_n_*) via time delays *τ_k_* of *V*(*t_n_*) [37, 1, 20] (*τ*_*n*+1_ > *τ_n_*):

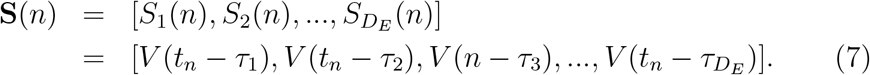

The use of Takens’ theorem in nonlinear dynamics [37] is widely practiced in the analysis of time series from nonlinear systems [1, 20].

The Physics behind the time delay construction is that as the the observed system moves from time *t_n_* – *τ*_*n*+1_ → *t_n_* – *τ_n_*, the dynamics of the system incorporates information about the activity of all other variables beyond the voltage alone. When the quantity *V*(*t_n_* – *τ_n_*) is approximately statistically independent of the quantity *V*(*t_n_* – *τ*_*n*+1_), each can be used as the components in a ‘proxy vector’ **S**(*n*) representing the system dynamics as it develops in dimensions higher than V(t) alone.

Using the average mutual information [12, 1, 20] between *S_j_* and *S_k≠j_*, and choosing time lags {*τ_j_*, *τ_k_*} giving a minimum of this average mutual information, we achieve an approximate information theoretic, nonlinear independence of the *D_E_* components of **S**(*n*) with respect to each other.

In principle in the discussion of Takens’ work, if one has an infinite amount of noise free data, any time delay or set of time delays [14] would give a proxy state vector **S**(*n*) that is equivalent to the original dynamical space of the data source. Of course, we never have that, so a ‘guide’ was devised by [12] which suggests that choosing the components of **S**(*n*) to be nonlinearly independent of each other, using average mutual information as a ‘nonlinear correlation’ among the components, would provide a good measure of the ability of the components of **S**(*n*) not to be parallel to each other. If that is achieved, then they would span the *D_E_* dimensional space of **S**(*n*) in a numerically useful manner. If there are multiple times scales in the data, the method of [14] could be a useful method to implement.

To estimate *D_E_* one may use the method of false nearest neighbors [1, 20].

It is typical, but not required, to select *τ_a_* = (*a* – 1)*τ*; *a* = 1, 2, … *D_E_*. In this standard choice of delay coordinates there are two parameters that determine the vectors **S**(*t*) : {*τ, D_E_*}. *τ* is conveniently taken to be an integer multiple of h; *D_E_* is an integer about twice the dimension of the system generating V(t). More precise results for these criteria are given in [31, 1, 20].

In **S** space we develop a map from time *t_k_* – *τ_k_* to time *t_k_* + *τ_k_* + *h*:

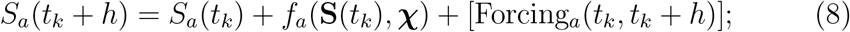

*a* = 0, 1, … *D_E_* We are interested in making steps of size h to arrive at the dynamical discrete time map (*a* = 1, 2, .., *D_E_*):

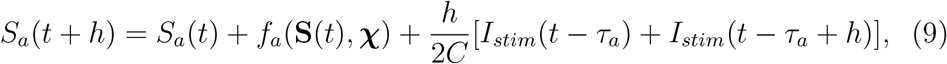

as each component of **S** is a voltage. The last term in Eq. (9) is the trapezoidal approximation to the integral of *I_stim_*(*t*) over the interval [*t_k_* – *τ_a_*, *t_k_* – *τ_a_* + *h*]. Each *f_a_*(**S**, **χ**) is a function of the *D_E_*-dimensional variables **S** and constants **χ** to be trained as before. The parameters are now **χ** = {*w_aq_*, *σ*, *β*, *τ*, *D_E_*}. We represent *f_a_*(**S**, **χ**) using a linear combination of Gaussian RBFs.

In Eq. (9) we have introduced *D_E_* vector fields *f_a_*(**S**, **χ**) whose parameters **χ** must be estimated by requiring Eq. (9) to be true over a training set with {*t_n_*}; *n* = 1, 2, …, *N* – 1. The trained dynamical map **S**(*n*) → **S**(*n* + 1) is used to forecast all components of **S**(*k*) for *k* ≥ *N*.

Since we are representing the development of voltage in each component of *f_a_*(**S**, **χ**) these vector fields should be independent of a, and, in practice, we take that as given and move forward only the component *S*_1_(*n*), namely the observed voltage, so

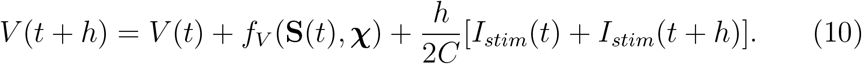

We then use that result to evaluate the remaining components of **S**(*n*) required in Eq. (9).

## 6 Results When Only V(t) is Observed

### 6.1 DDF Analysis of Numerical V(t) Data from an NaKL Neuron

Still using data from the numerical solution of the basic HH equation, Eq. (6), we now train a Gaussian RBF via Eq. (9). 125ms of data for V(t) alone are employed in the estimation of the parameters **χ** of the RBF.

The trained DDF is used to predict the subsequent 500ms of the V(t) time course.

As in the earlier (unrealistic) example when all state variables from the NaKL neuron model were available, when V(t) alone is presented, the DDF neuron is able to predict the time course of the observable membrane voltage with significant accuracy. This result is shown in Fig. (2).

**Figure 2:**
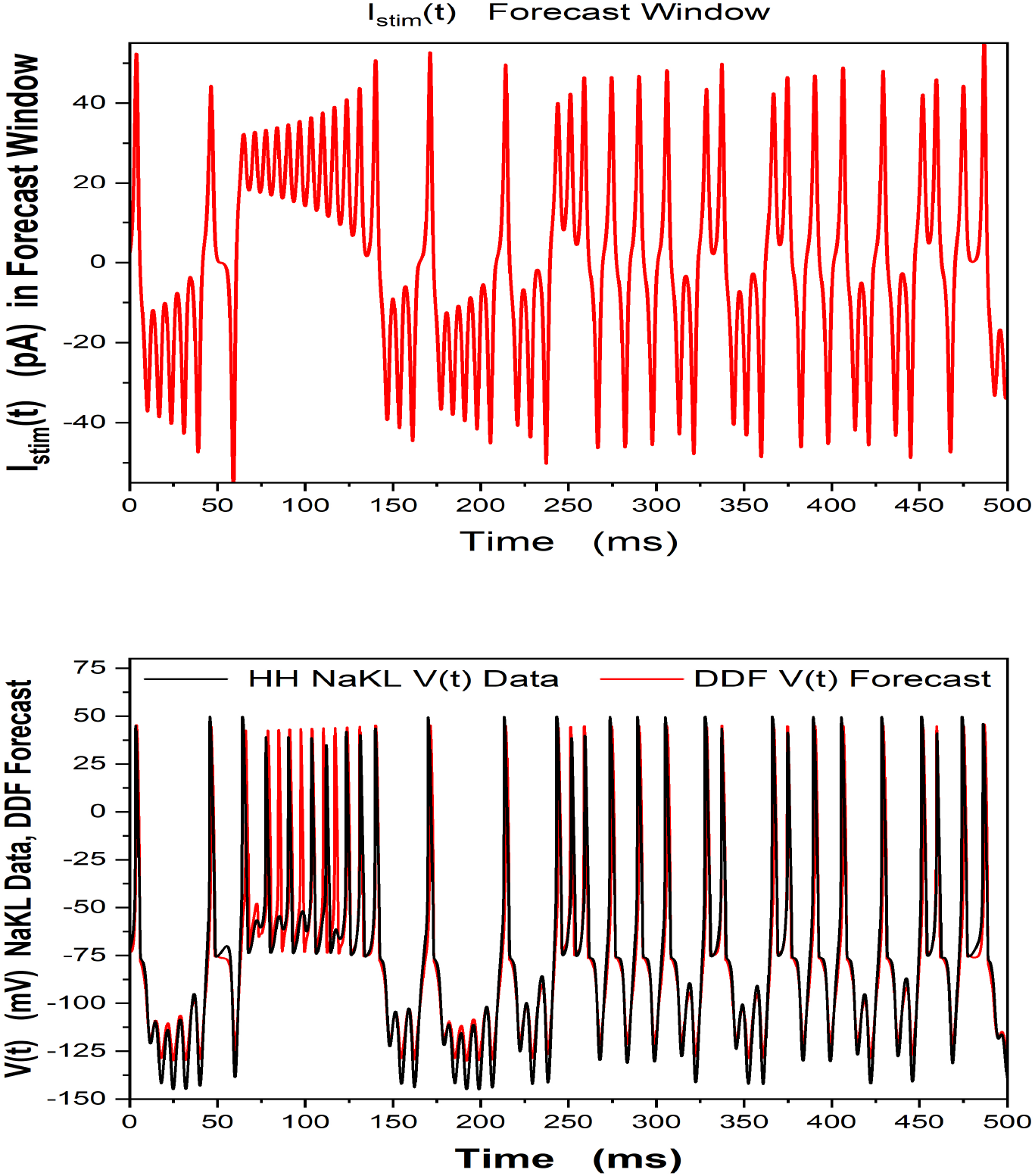
Data is generated by solving the HH-NaKL model Eq. (6). We observe only the membrane voltage, V(t), and use 125ms of these data for training the Gaussian RBF. **Top Panel** *I_stim_*(*t*) in the 500ms forecasting window. **Bottom Panel** We display the V(t) forecast of the trained DDF-NaKL construct for 500ms after the training window and compare it to the HH NaKL model generated V(t) data. `This is a numerical calculation, but it corresponds to a realistic current clamp experiment where, given a driving current` *I_stim_*(*t*), `only V(t) is observed`. h = 5 × 10^−3^ms, *β* = 100, R = 10^−3^, *τ* = 8*h*, *N_c_* = 5000.

Note that one cannot forecast the gating variables {**A**(*t*)} of the HH NaKL model as we have no observed information about them. Through the time delay vector **S**(*t_n_*) the biophysical information in the {**A**(*t*)} is encoded in the trained parameters **χ**.

#### 6.1.1 Comparing DDF Forecasting and NaKL Forecasting Times

To assess the effectiveness of using a trained DDF to forecast V(t) data, for example for the efficiency of computational demands on a DDF neuron in a circuit where it replaces a HH model, we compared the computation time for solving our HH NaKL model to the forecasting time of a DDF trained on the V(t) from that HH NaKL model.

We generated HH NaKL data by solving Eq. (6) using a standard fourth order Runge-Kutta method [29, 28] with a time step of h = 0.02 ms. The times taken by the generation of the NaKL data in a forecast window of 2000 ms (10^5^ time steps) were 8.9 s for either CPU time or wall clock time.

We then forecast in the same window using the same *I_stim_*(*t*) as for the HH NaKL model but using a trained DDF, trained on V(t) from the HH NaKL model and forecasting only V(t).

The choice of the number of centers *N_c_* in the training and forecasting for the DDF is important to the forecasting time of the DDF. If we choose *N_c_* = 500, then the CPU time for forecasting the 2000 ms with h = 0.02 is 3 s, while the wall clock time is 2.4 s. If we decrease the number of centers to *N_c_* = 100, the the CPU time during forecasting is reduced to 2s while the wall clock time drops to 1.5 s.

The training time for the DDF with *N_c_* = 500, on V(t) from the HH NaKL models is 1.1s for CPU time and 0.63 s for wall clock time. This decreases to 135 ms CPU time and 142 ms wall clock time for *N_c_* = 100.

These times will vary as the complexity of the HH model neuron increases from the minimalist NaKL model to a model for observed laboratory observations. One expects the V(t) trained and forecasting DDF to become relatively more efficient than our results on the simple NaKL model. The training times for a DDF on V(t) data alone are quite fast. In the scenarios where we substitute DDF V(t) neurons for HH neurons in a circuit the computational efficiency is what will be of central importance.

### 6.2 DDF Analysis of Laboratory Data from an HVC Neuron in the Avian Song System

With these DDF results on numerical data generated by the solution of the NaKL HH equation in hand, we turn to the use of DDF when presented with experimental current clamp data.

In Fig. (3) we show the stimulating current *I_stim_*(*t*) (**Left Panel**) and the resulting membrane voltage time course (**Right Panel**) from an *in vitro* current clamp experiment on an isolated neuron in the HVC nucleus of the zebra finch song system [27].

**Figure 3:**
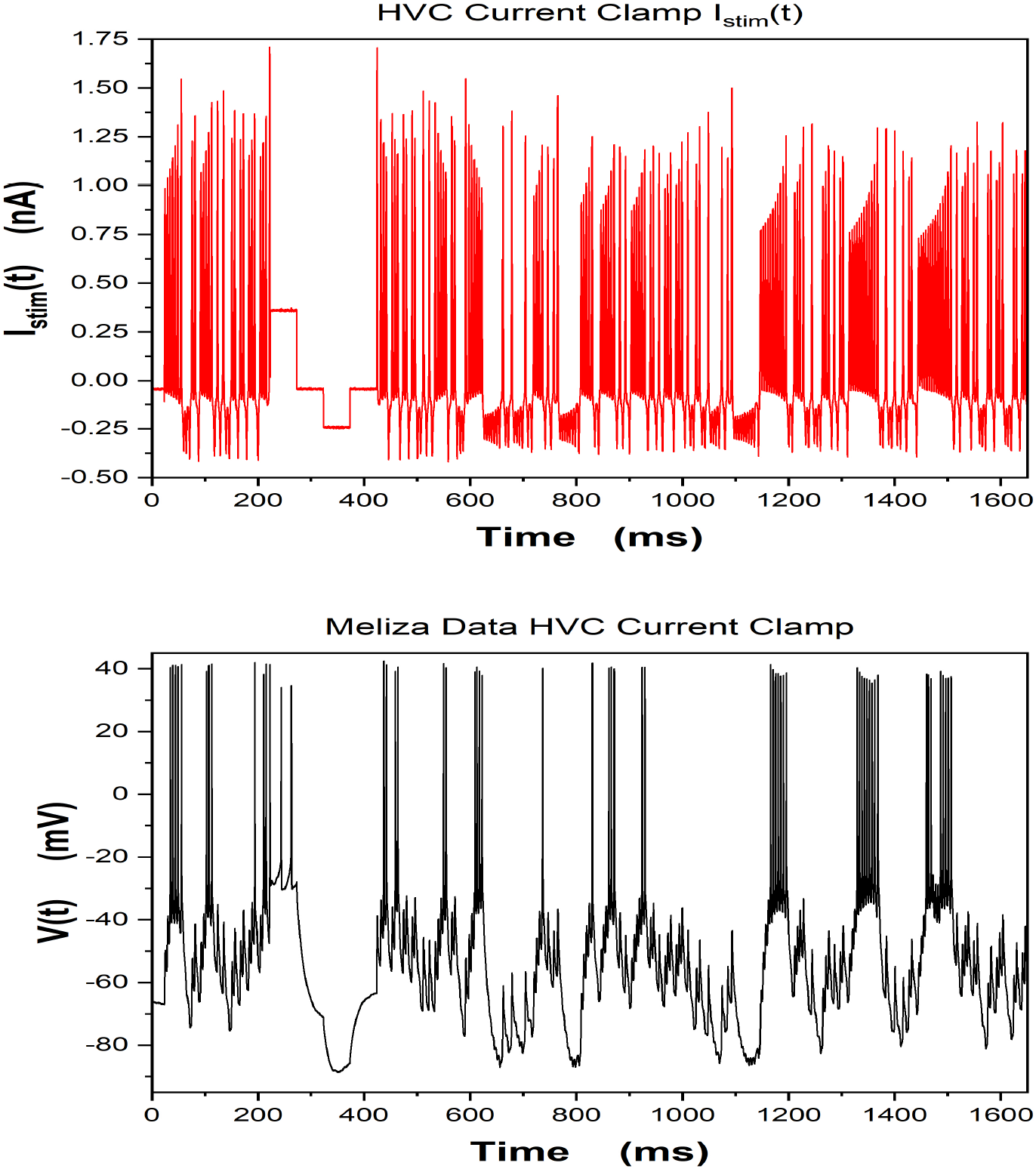
**Top Panel** The stimulating current *I_stim_*(*t*) presented to an isolated neuron in the HVC nucleus of the zebra finch song system in an *in vitro* current clamp experiment at the University of Chicago laboratory of Dan Margoliash. **Bottom Panel** The recorded membrane voltage response to *I_stim_*(*t*). These data were collected by C. D. Meliza (now at the University of Virginia) who designed *I_stim_*(*t*) in collaboration with M. Kostuk, then a UCSD Physics PhD student.

### 6.3 Only V(t) is Observed in Laboratory Current Clamp Data

Current clamp data was collected by C. D. Meliza at the University of Chicago laboratory of Daniel Margoliash from presenting various stimulating currents *I_stim_*(*t*) to isolated HVC neurons in a zebra finch *in vitro* preparation. The data were organized into ‘Epochs’ of length about 2-6 seconds observed over several hours.

In Fig. (4) we show the results of constructing a DDF neuron forecasting model on these data. The first 500ms of the stimulating current data and the V(t) response data (these are not shown) were used to train a DDF RBF model. In the **Left Panel** we show the stimulating current used in 500ms of a prediction window for the same experimental preparation. In the **Right panel** we show the voltage forecast of V(t) using the trained DDF model (blue) along with the observed voltage response (black).

**Figure 4:**
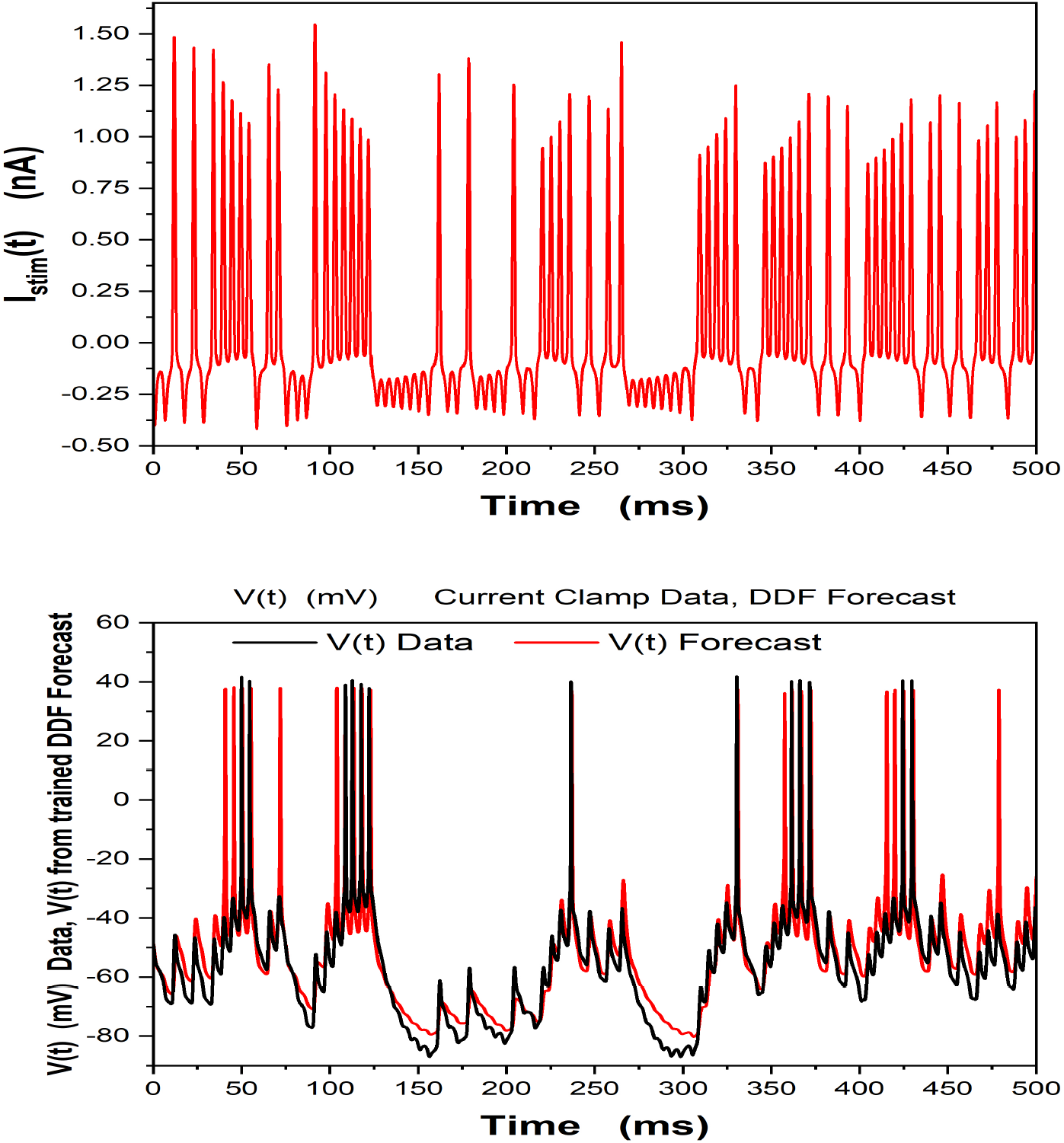
A Gaussian RBF representation for the vector field of the membrane potential dynamics was used to train a DDF. The training used 500ms of observed V(t) data. **Top Panel** In the 500ms forecast window *I_stim_*(*t*) data from Fig. (3) were used. **Bottom Panel** The V(t) forecast for 500ms by the trained DDF neuron (in red) along with 500ms of the observed V(t) current clamp data (in black). h = 0.02 ms, *τ* = 2*h*, *N_c_* = 5000, *D_E_* = 3.

Next we wish to provide the same analysis as in Fig. (4) using two different epochs, Epoch 25 and Epoch 26, of data collected by C. D. Meliza in the Margoliash laboratory. In Fig. (5) we display the stimulating current *I_stim_*(*t*) used in Epoch 25, **Upper Panel**, and the V(t) response of the neuron in the **Bottom Panel**. In Fig. (6) we display the stimulating current *I_stim_*(*t*) used in Epoch 26, **Upper Panel**, and the V(t) response of the neuron in the **Bottom Panel**. This demonstrates that the DDF model neuron, just as the HH model neuron, responds appropriately to changes in the stimulating current. It is the unknown intrinsic properties of the neuron for which we have introduced a RBF representation.

**Figure 5:**
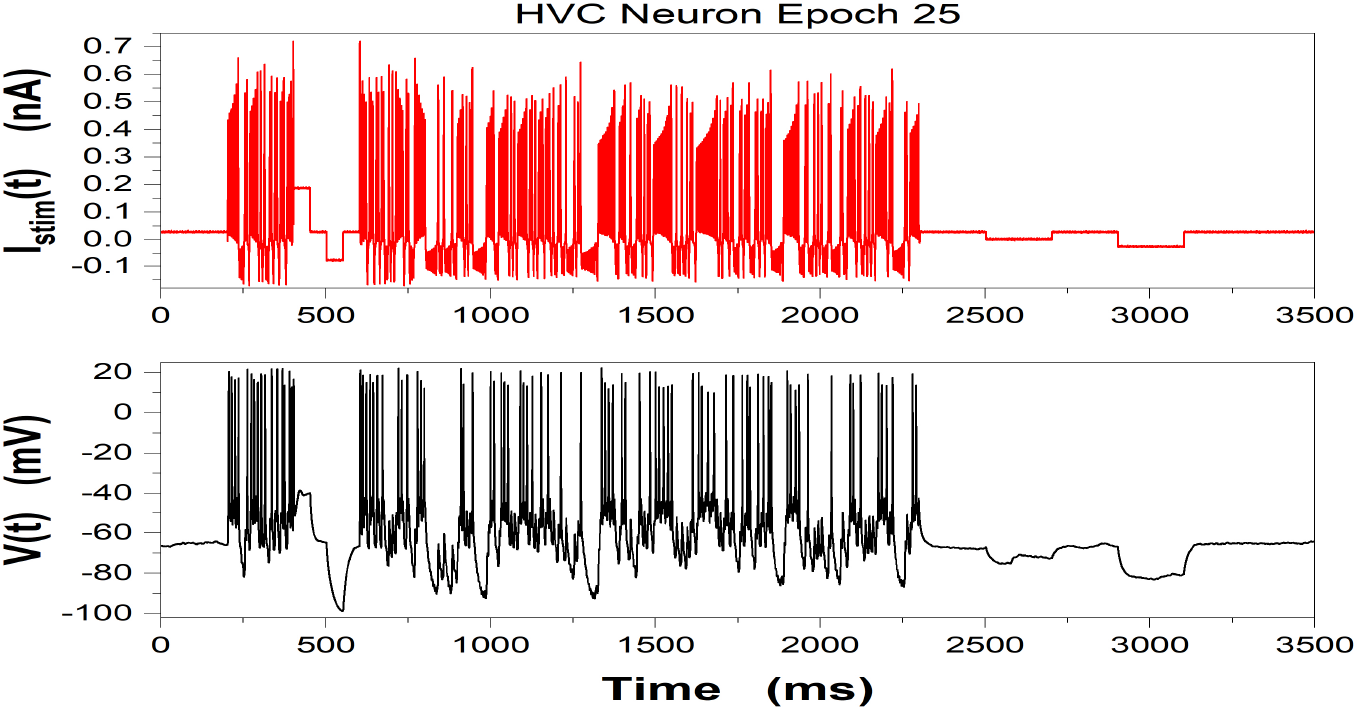
**Top Panel** Stimulating current *I_stim_*(*t*) for a current clamp experiment at the Margoliash laboratory of the University of Chicago. **Bottom Panel** Membrane voltage response, V(t), to *I_stim_*(*t*). Data was collected by C. D. Meliza (now at the University of Virginia) in sequential time ‘epochs’ from the same HVC neuron in Zebra Finch. Between epochs *I_stim_*(*t*) = 0. Many epochs of varying length in time and with different stimulating currents *I_stim_*(*t*) were recorded. These data are 3500ms from Epoch 25 of the observations.

**Figure 6:**
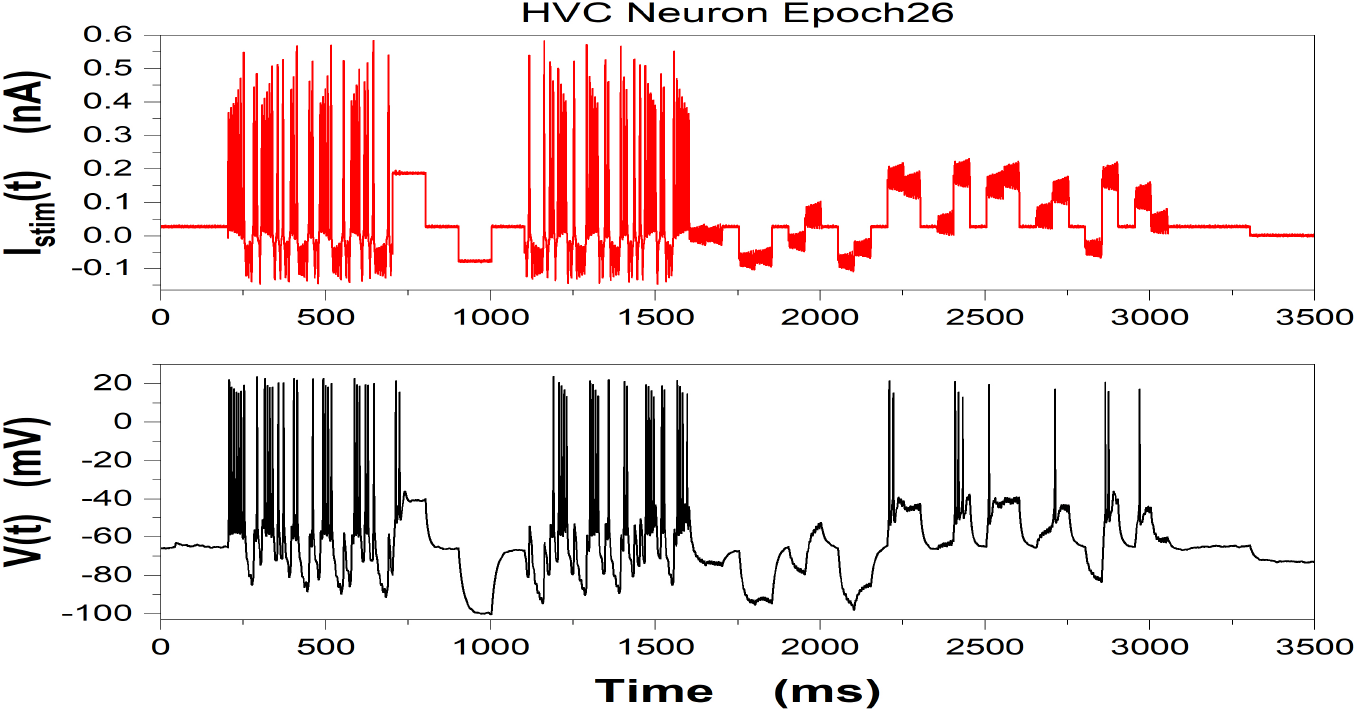
**Top Panel** Stimulating current *I_stim_*(*t*) for a current clamp experiment at the Margoliash laboratory of the University of Chicago. **Bottom Panel** Membrane voltage response, V(t), to *I_stim_*(*t*). Data was collected by C. D. Meliza (now at the University of Virginia) in sequential time epochs from the same HVC neuron in Zebra Finch. These data are 3500ms from Epoch 26 of the observations.

### 6.4 Training a DDF Neuron in One Epoch and Using it to Forecast in Another Epoch on Experimental Current Clamp Data

Another informative test of the DDF approach is to train a DDF neuron on neuron data from one time epoch with a selected *I_stim_*(*t*), then using the same parameters **χ** from the first epoch forecast the V(t) response to different *I_stim_*(*t*) presented in a second time epoch. This a test of the DDF neuron ability to correctly respond to different stimulating currents.

In Fig. (5), (**Top Panel**), we show *I_stim_*(*t*) in Epoch 25 and the resulting, *V_Data_*(*t*) (**Bottom Panel**). In Fig. (6), (**Top Panel**), we show *I_stim_*(*t*) in Epoch 26 and the resulting, *V_Data_*(*t*) (**Bottom Panel**). These data are from two epochs of a current clamp experiment from the Margoliash laboratory (University of Chicago) on an isolated neuron from the zebra finch HVC nucleus.

Next, in Fig. (7) we display the observed data, then we analyze the ability of DDF trained on data from Epoch 25 to forecast within that Epoch. Then in Fig. (8) we show how the DDF model trained on data from Epoch 25 is able to forecast the V(t) for data in Epoch 26 where the *I_stim_*(*t*) is different, though the neuron is the same.

**Figure 7:**
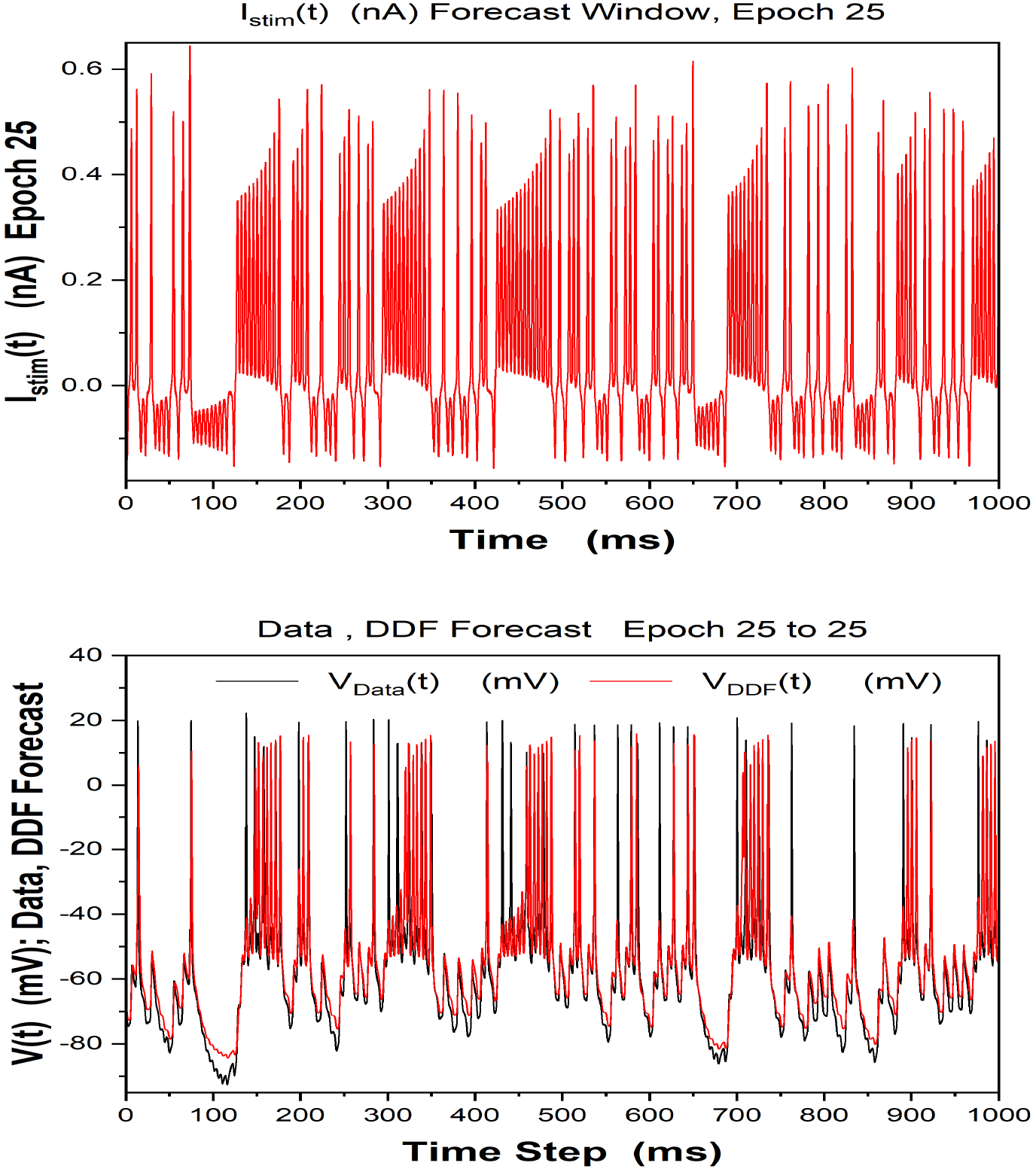
DDF forecasting and Observed Data; Epoch 25. Training window was 1000ms. Only V(t) was observed and used to train the DDF neuron. **Top Panel** Observed *I_stim_*(*t*) in Forecast Window **Bottom Panel** Forecast for 1000ms by DDF Voltage on Epoch 25 data. h = 0.02 ms, *τ* = 2*h*, *N_c_* = 5000, *D_E_* = 4, R = 10^−3^, *β* = 10^−3^.

**Figure 8:**
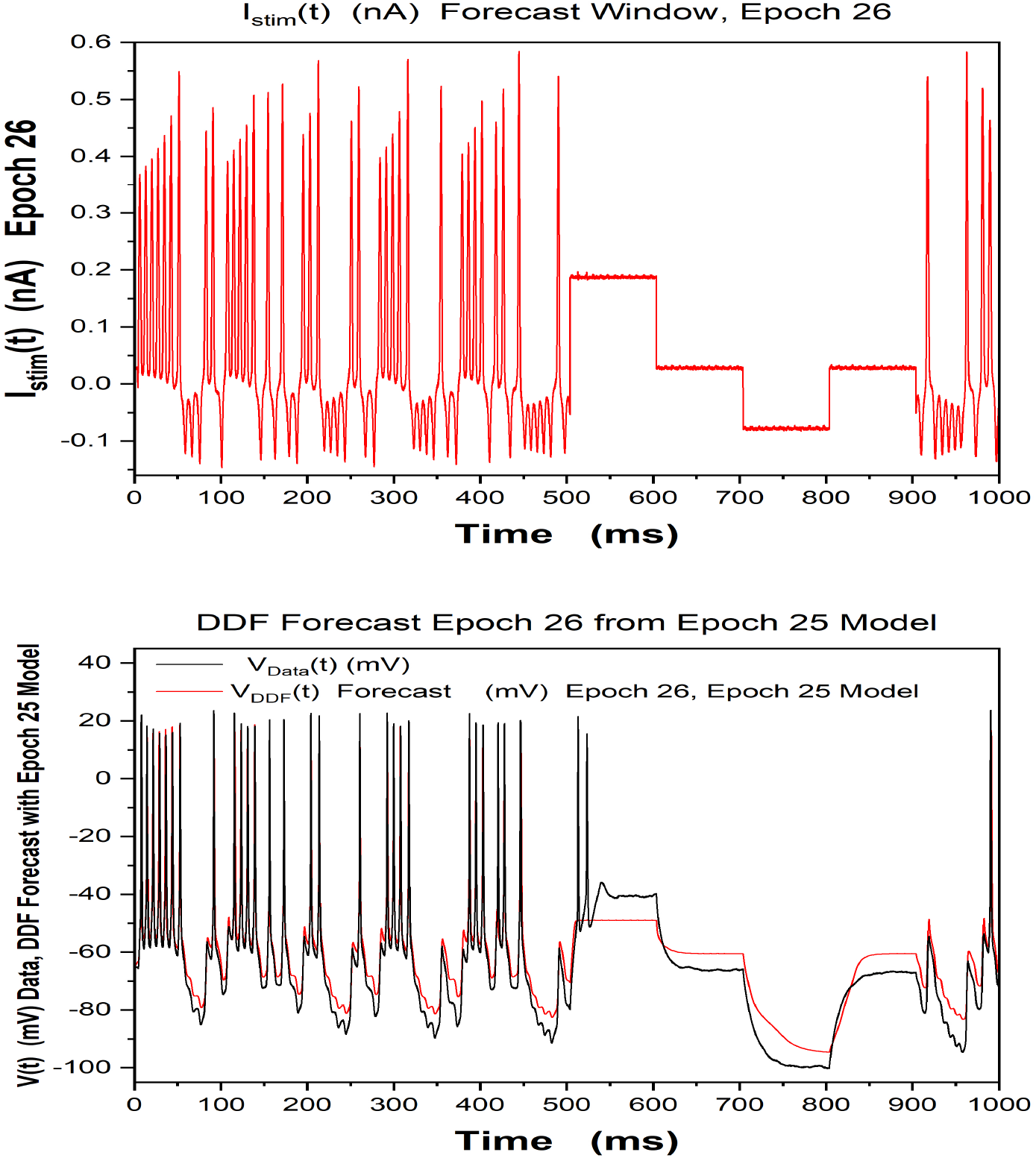
Analysis of Epoch 26 data using a DDF neuron trained on 1000ms Epoch 25 V(t) data. **Top Panel** *I_stim_*(*t*) for Epoch 26 in the Forecast Window. This is different from the *I_stim_*(*t*) used in the Epoch 25 training window. **Bottom Panel** Observed current clamp Data and DDF forecast in the Epoch 26 time window using the Epoch 25 trained DDF. The performance is worst for regions of *I_stim_*(*t*) comprised of square pulses; this is consistent with the observations in [24]. *h* = 0.02 ms, *τ* = 2*h*, *N_c_* = 5000, *D_E_* = 4, R = 10^−3^, *β* = 10^−3^.

### 6.5 Comments on the HVC Current Clamp Experiments

There is a large library of current clamp data from this preparation. The observation window for the data used here was about 1650-3500ms. Many entries in the library have a longer window of time over which data was collected, and this would allow longer training windows to be used. In previous work with this kind of data [21, 27] longer estimation windows typically result in better forecasting.

The stimulating currents in these data were designed with three biophysical principles in mind: (a) the amplitude variations of *I_stim_*(*t*) must be large enough to generate many action potentials as well as substantial periods of sub-threshold behavior. This guarantees that the full dynamic range of neuron response is well represented. (b) The observation window must be long enough in time to assure the same goals as in (a). (c) The frequencies in *I_stim_*(*t*) should be low enough so that properties of the stimulating signal are not filtered out by the RC low-pass characteristic of the cell membrane. If these criteria are not met, aspects of *I_stim_*(*t*) are filtered out by the cell membrane and the training is likely to be insufficiently well informed. V(t) data collected with *I_stim_*(*t*) chosen employing these guidlines were regularly successful in using DA to estimate the properties of rich HH models [38, 21, 27] from laboatory data.

The context of this discussion is expanded in Appendix C.

## 7 Adding Other Observables

In many neurobiological investigations more observables than just V(t) may be available. We examine how these may be combined with observations of V(t) in a DDF framework or used on their own.

As an example, we consider the important quantity of the time course of the intracellular concentration of calcium [*Ca*^2+^](*t*) = *Ca*(*t*). This is governed by a conservation equation of the form

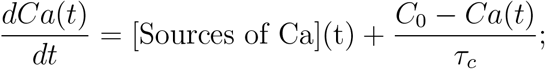

where *C*_0_ is an equilibrium or rest state concentration of Ca, and *τ_c_* Is a relaxation time for the *Ca*(*t*) dynamics.

The equation for the difference Δ(*t*) = *Ca*(*t*) – *C*_0_, is

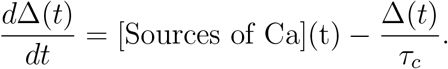

The sources of Ca ions are attributed to voltage gated Ca channels with various properties and release of and uptake of Ca from intracellular stores such as the Endoplasmic Reticulum; [16, 17, 11, 41].

Integrate Eq. (11) from time t to time t + h to make a discrete time map

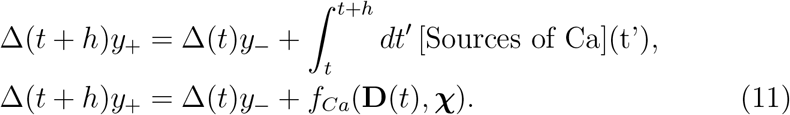

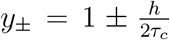 appears here as we identify the appearance of Δ(*n*) and Δ(*n* + 1) on both sides of the equation for the flow of *Ca*(*t*). The integral over Δ(*t*) uses the trapezoidal approximation and the unknown dynamics for the sources and sinks of *Ca*(*t*) are represented within the RBF vector field *f_Ca_*(**D**(*t*). **χ**).

The *Ca*(*t*) time variation is a projection from higher dimensional dynamics of a neuron and a time delay ‘unprojection’ is required here as well. The time delay state vector for this situation is

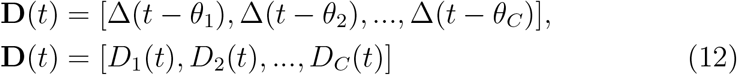

in which the time delays {*θ_k_*}; *k* = 1, 2, …, *D_C_* appear. This construct ‘unprojects’ the projected observation of *Ca*(*t*).

If V(t) and *Ca*(*t*) were both to be observed, then the ‘unprojection’ occurs in the joint time delay space of voltage, **S**(*t*), Eq. (7), and **D**(*t*), Eq. (12). Further observations, when available, may be added to this framework.

## 8 Using DDF Neurons in a Network

One important goal of using neuron models trained by data alone, i.e. DDF neurons, is to provide a reduced model based on biophysical observations to employ in building network models.

We demonstrate this in the most basic network comprised of just two neurons, connected by gap junctions, as shown in Fig. (9).

**Figure 9:**
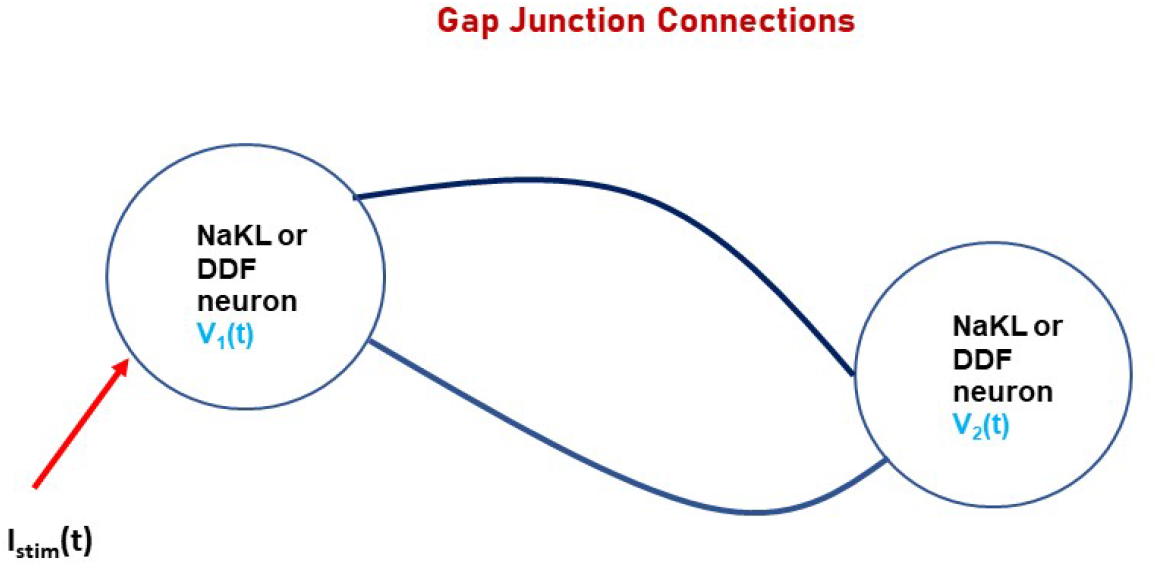
A two neuron circuit comprised of two NaKL HH neurons **or** two DDF neurons trained on NaKL voltage data. There are gap junction connections between the two neurons in the circuit. The circuit is driven by the stimulating current I*_stim_*(*t*) presented to neuron one. An NaKL neuron is a Hodgkin-Huxley model neuron with Na, K, and leak channels [18, 34]. The DDF neuron is one built with RBFs trained with V(t) data from the HH NaKL model. The computational task using the DDF neurons is substantially simplified as only membrane voltage plays a role and no integration of HH differential equations is required in establishing the behavior of the neural circuit. The equations of the map are given in Eqs. (14), (15), and (16)

Note that it is only the presynaptic and postsynaptic voltages that convey information from any neuron in this circuit to others in the circuit. Either the HH model NaKL neuron or the NaKL trained DDF neuron may be used in this small network. Each produces voltage signals that couple the neurons.

### 8.1 Discrete Time Gap Junction Dynamics

The differential equations for the two neuron circuit with gap junction connections are these:

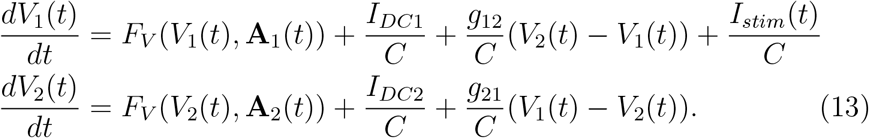

Integrating these over the interval [*t, t* + *h*], we arrive at

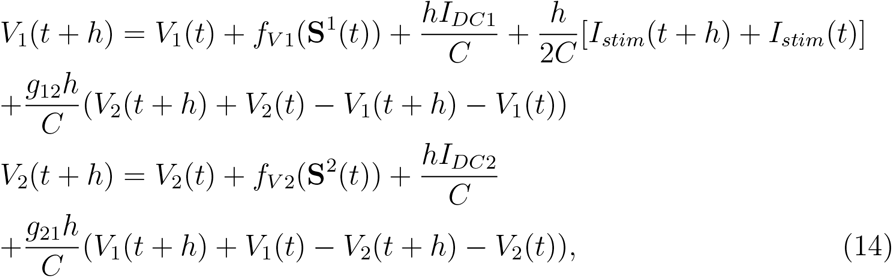

and then

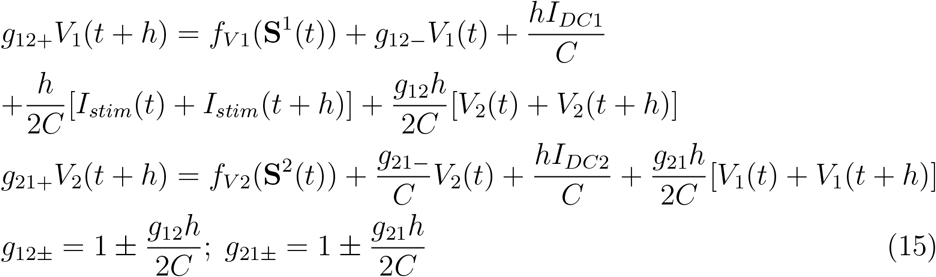

Defining the two dimensional vector **v**(*t*) = [*V*_1_(*t*), *V*(*t*)] Eq. (15) may be put into matrix form

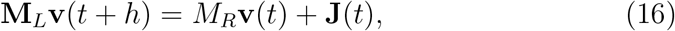

in which

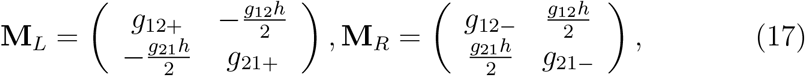

and 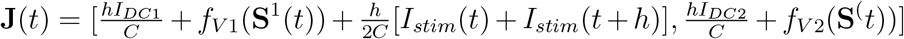.

The desired discrete time map for gap junction coupling in a two neuron map is then

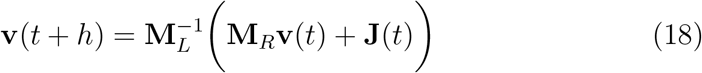

## 9 Dynamics of A Simple Two Neuron Network

We are now prepared to use the dynamical discrete time maps for circuits such as the one in Fig (9).

This proceeds as follows:

a. determine the neuron RBFs for the two neurons *f*_*V*1_(**S**^1^) and *f*_*V*2_(**S**^2^). In the simplest case, which we adopt here, the neurons are the same, and the RBFs, *f_V_*(**S**), are the same function of their respective multivariate arguments **S**^1^(*t*) and **S**^2^(*t*).
b. select a stimulating current *I_stim_*(*t*),
c. select excitatory or inhibitory synaptic connections or gap junction connections,
d. using the DDF neurons in place of the HH neurons appearing in the circuit use the trained DDF neurons, and
e. evaluate the network behavior using the discrete time map for the coupled *V*_1_(*t_n_*) and *V*_2_(*t_n_*), Eqs. (14) and (15), when DDF neurons are at the nodes of the network.

### 9.1 A Two Neuron Circuit: HH NaKL Neurons or DDF, NaKL Trained, Neurons; Gap Junction Connections

We begin by using a selected *I_stim_*(*t*) to an HH-NaKL neuron and use the resulting V(t) data to train a DDF discrete time map for V(t) generated from an HH NaKL neuron. The forecasting ability of the DDF is shown in Fig. (10). We now have the HH model NaKL and the DDF model NaKL neuron we require for a comparison of the circuit using one and then the other at the nodes of the two neuron network.

**Figure 10:**
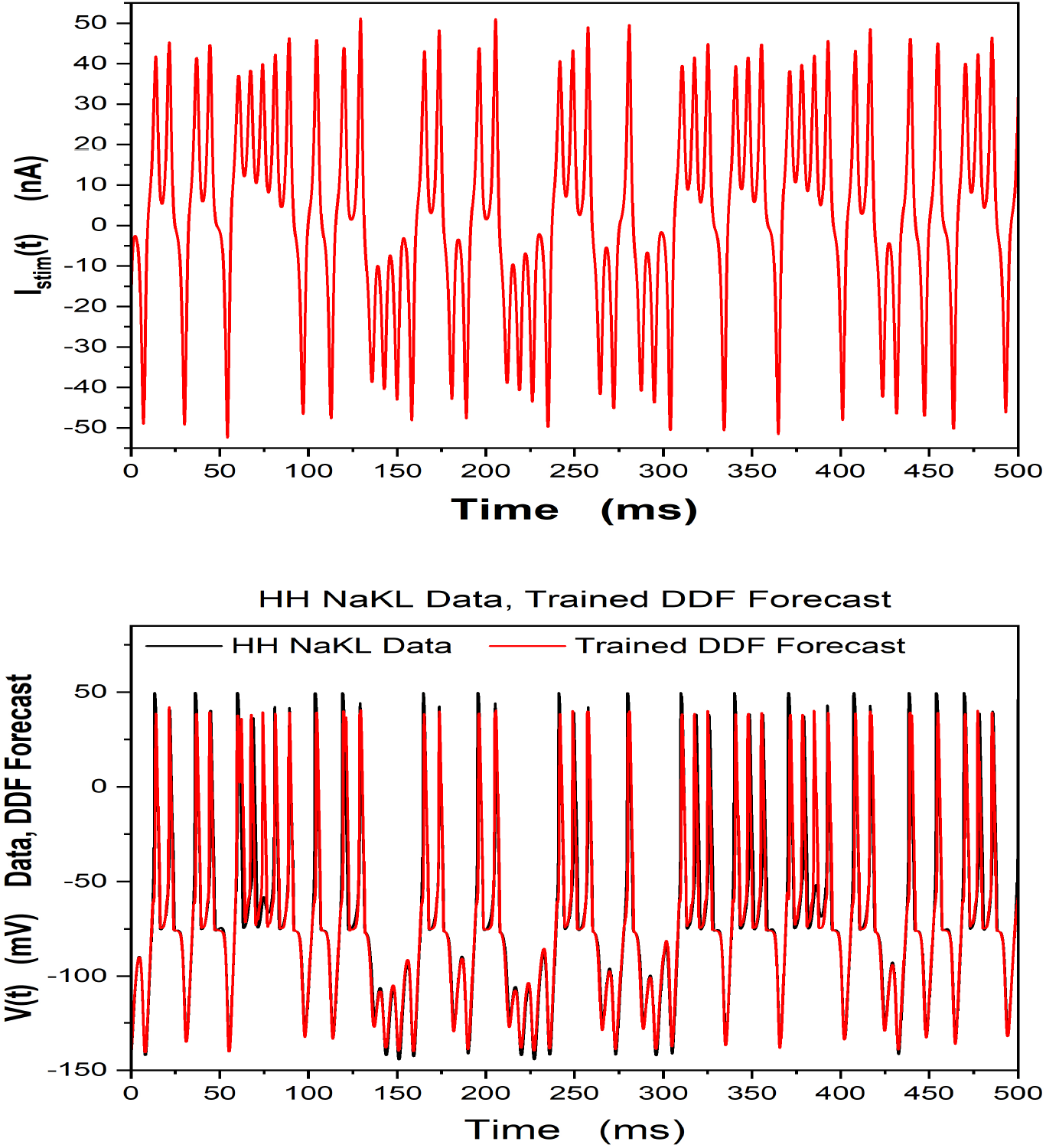
An HH-NaKL model neuron and a DDF-NaKL were driven by the same *I_stim_*(*t*) in a training window of 500ms. Only the V(t) data from the HH-NaKL model neuron was used to train the DDF-NaKL neuron. (h = 0.02ms, *D_E_* =3, *β* = 10, *τ* = 3h, *N_c_* = 5000.) In the **Top Panel** we show the *I_stim_*(*t*) in a subsequent 500ms window used for prediction by the trained DDF-NaKL model. **Bottom Panel** Comparison of the voltage time courses V(t) from the numerical solution of the HH-NaKL and the forecast of the trained DDF-NaKL models over a 500ms forecasting window. This V(t) trained DDF-NaKL neuron was used in the 2 neuron gap junction network, Fig. (9), to replace the HH NaKL neuron.

Using the HH-NaKL neuron at both nodes of the network Fig. (9) we generated the time course of *V*_1_(*t*) and *V*_2_(*t*). Then replacing the HH-NaKL neurons with the trained DDF-NaKL neuron we generated another set of *V*_1_(*t*) and *V*_2_(*t*) time courses.

A comparison of results of using these two neuron models at the nodes of our simple network is shown in Fig. (11). While the ‘network’ we selected is simple, the idea that we are able to replace an HH-neuron with a trained DDF-neuron in a network is now supported by these results..

**Figure 11:**
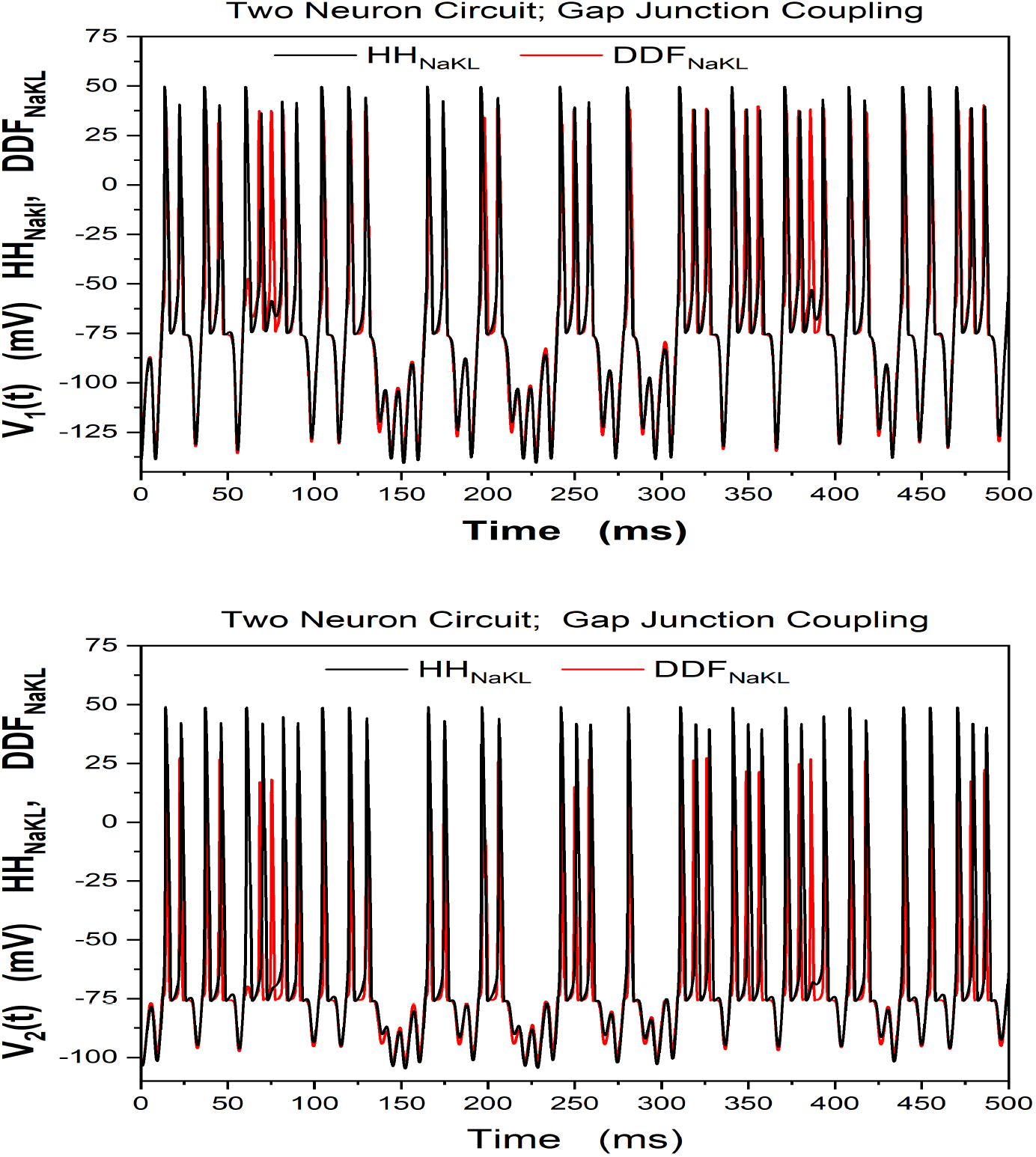
In the two neuron circuit with gap junction connections shown in Fig.(9) HH-NaKL model neurons were used in the circuit and *V*_1_(*t*) and *V*_2_(*t*) were recorded. Then we used the trained DDF-NaKL neurons, see Fig (10) in the same circuit with the same *I_stim_*(*t*). *V*_1_(*t*) and *V*_2_(*t*) were recorded with HH-NaKL neurons and with DDF neurons. **Top Panel**. Comparison of the time courses of *V*_1_(*t*) in the two circuits. **Bottom Panel** Comparison of the voltage time courses *V*_2_(*t*) in the two circuits.

This result is, as noted, for gap junction couplings between the two neurons. The way one introduces ligand gated synaptic connections into a network is discussed in Appendix B.

### 9.2 One DDF Neuron Driving a Second DDF Neuron through a Synaptic Connection

To explore the ability of DDF neurons to work in a biological network with ligand gated synaptic connections, we constructed a network segment in which Neuron 1, with membrane voltage *V*_1_(*t*), is driven by a stimulating current *I_stim_*(*t*) and this neuron drives a second neuron, with membrane voltage *V*_2_(*t*), via an excitatory ligand gated synapse. This network segment is shown in Fig. (12).

**Figure 12:**
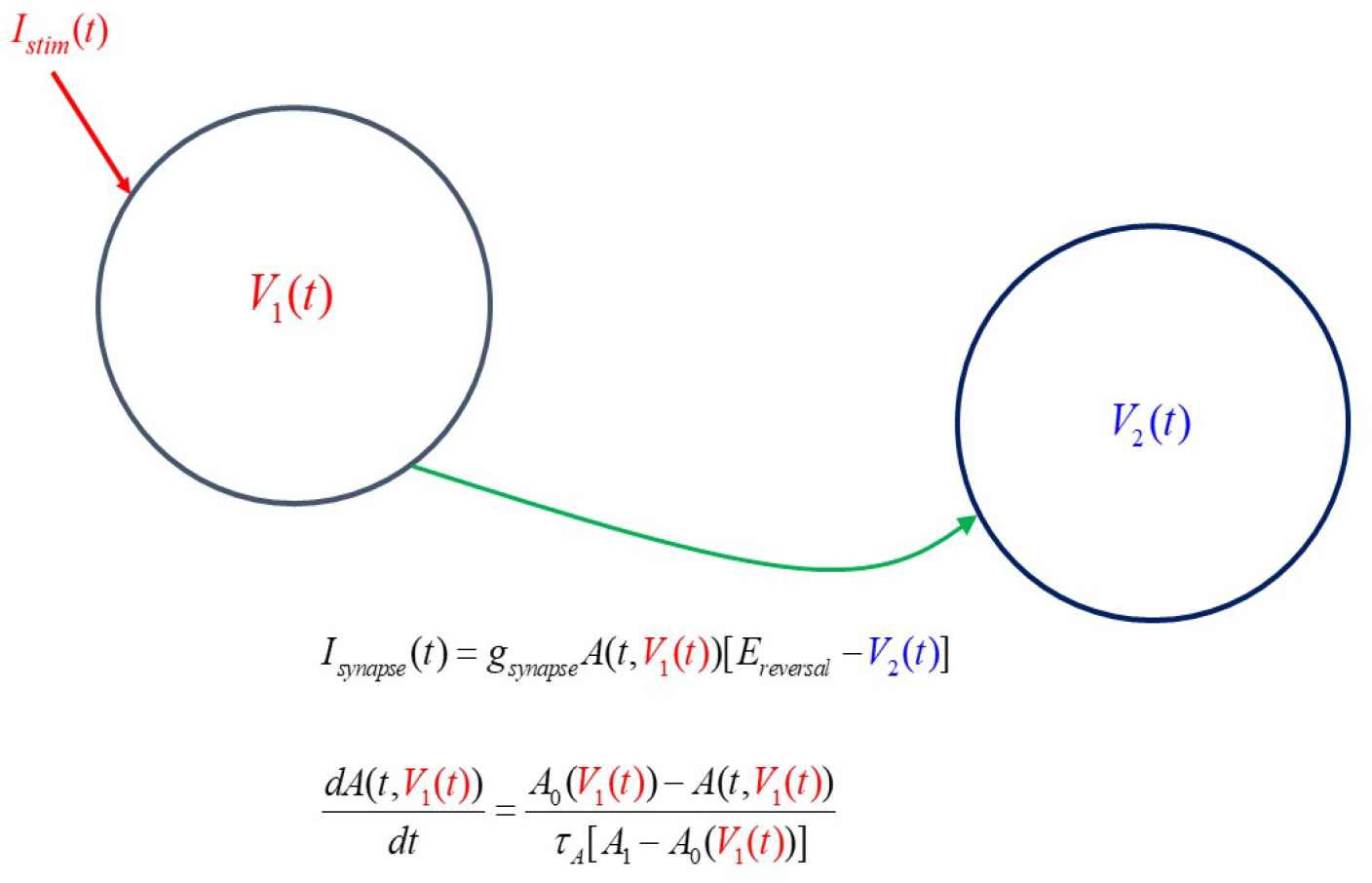
A network segment which has a presynaptic neuron with membrane voltage *V*_1_(*t*) driven by a stimulating current *I_stim_*(*t*) connected to a postsynaptic neuron with membrane voltage *V*_2_(*t*) by a ligand gated synaptic connection. The current flowing into neuron two is given in Eq. (19).

The synaptic current from the presynaptic neuron with voltage *V*_1_(*t*) and driving the postsynaptic neuron with voltage *V*_2_(*t*) is described by

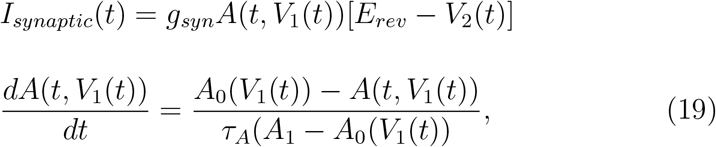

where *A*(*t*, *V*_1_(*t*)) is a synaptic gating variable. It is opened, *A*(*t*, *V*_1_(*t*)) ≈ 1, when neurotransmitter binds onto receptors on the postsynaptic cell. It is closed, *A*(*t*, *V*_1_(*t*)) ≈ 0, when that neurotransmitter is released from the postsynaptic receptor.

We represent driver of this transition in the neighborhood of a transition at a voltage *V*_0_ from closed to open by writing

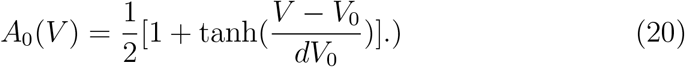

This function moves from very near 0 when *V* ≪ *V*_0_ to very near 1 when *V* ≫ *V*_0_, as desired, and it does so over an interval in voltage *dV*_0_.

For the excitatory synaptic connection, we selected *g_syn_* = 0.5, *E_rev_* = 50*mV*, *τ_A_* = 0.1*ms*, *A*_1_ = 9/8, *V*_0_ = –50*mV* and *dV*_0_ = 10*mV*.

The performance of this network segment was evaluated using the simple NaKL HH neuron as both the presynaptic neuron and the postsynaptic neuron. Along with the dynamics of the synaptic gating variable *A*(*t*, *V*(*t*)) this is a nine dimensional dynamical system. Selecting *I_stim_*(*t*) as we have earlier, we solved for *V*_1_(*t*) and *V*_2_(*t*) and stored these data for later use.

Next, using just the V(t) from an isolated NaKL neuron driven by the selected *I_stim_*(*t*) we built a DDF neuron using the time delay method described earlier. In the construction of the DDF neuron model we used *N_c_* = 500, *D_E_* = 4, *τ* = 2, and *h* = 0.02 ms in the data.

Two of these DDF neurons were then used in the network segment in place of the simple NaKL HH neurons, and the same network segment was driven by the same *I_stim_*(*t*) presented to neuron 1 connected by the same synaptic dynamics to DDF neuron 2. The DDF neurons were trained with 500 ms of HH model V(t) data.

In Fig. (13) we display the behavior of *V*_1_(*t*) from the presynaptic HH neuron and from the presynaptic DDF neuron when operating in the network segment.

**Figure 13:**
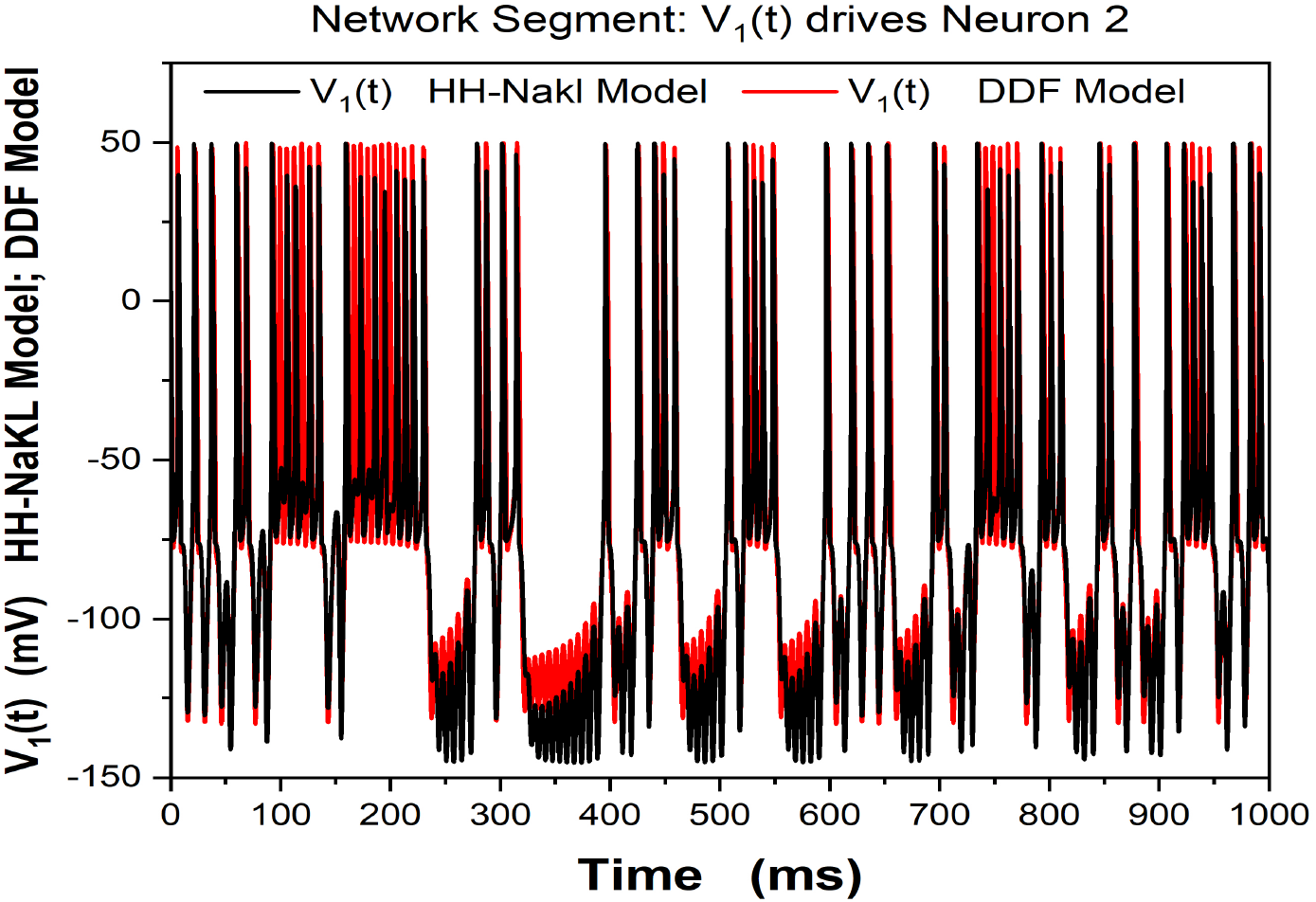
In the network segment shown in Fig. (12) Neuron 1 with membrane voltage *V*_1_(*t*) is driven by our selected *I_stim_*(*t*). The voltage activity *V*_1_(*t*) drives Neuron 2 through a ligand gated synapse, Eq. (19). In this figure we show *V*_1_(*t*) comparing *V*_1_(*t*) when HH NaKL neurons are in the network segment (black) and when DDF neurons, trained on the HH *V*_1_(*t*), are in the neuron segment (red). *N_c_* = 500 for the DDF neurons.

In Fig. (14) we display the behavior of *V*_2_(*t*) from the postsynaptic HH neuron and from the postsynaptic DDF neuron when operating in the network segment.

**Figure 14:**
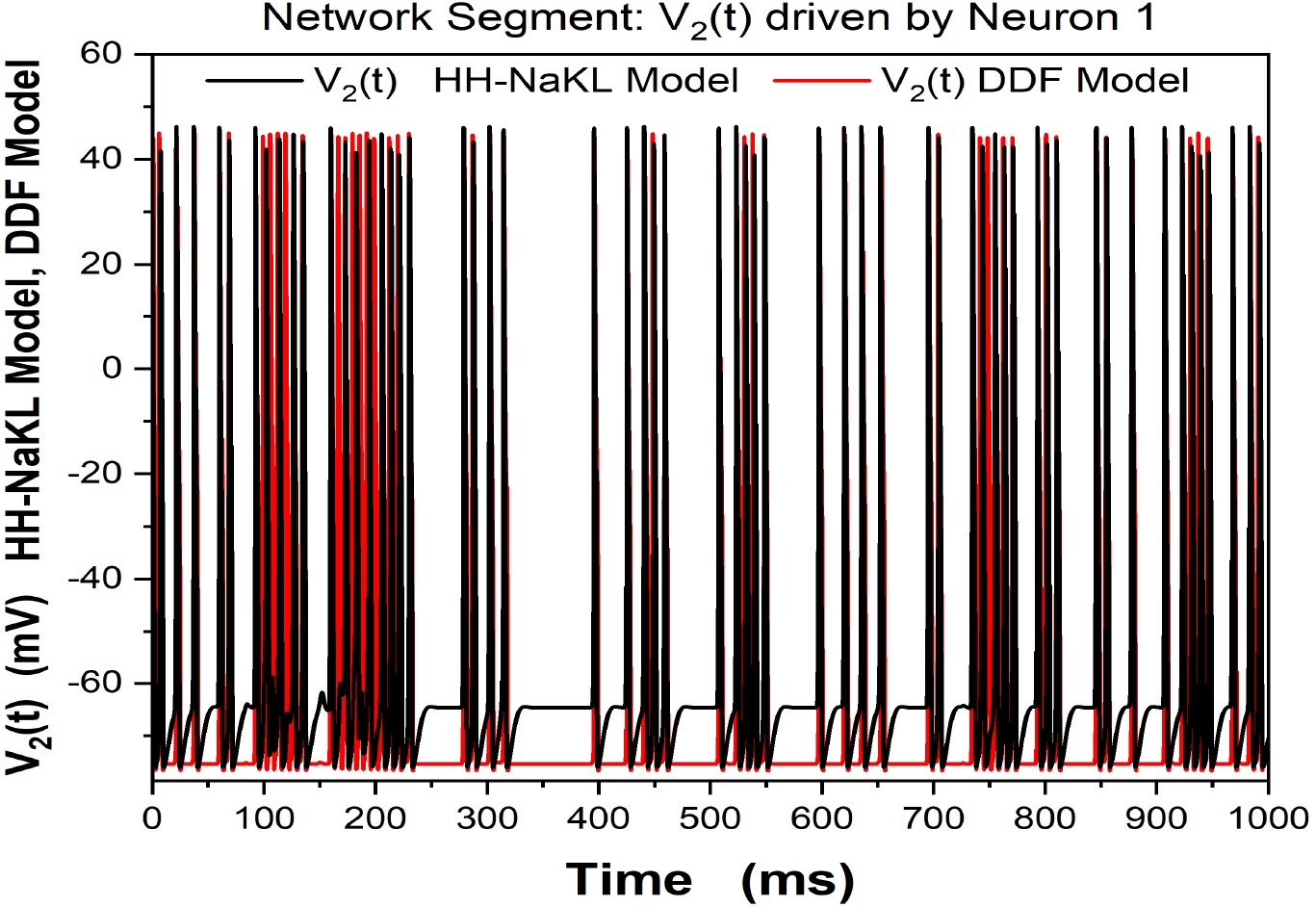
In the network segment shown in Fig. (12) Neuron 1 with membrane voltage *V*_1_(*t*) is driven by our selected *I_stim_*(*t*). The voltage activity *V*_1_(*t*) drives Neuron 2 through a ligand gated synapse, Eq. (19). In this figure we show *V*_2_(*t*) comparing *V*_2_(*t*) when HH NaKL neurons are in the network segment (black) and when DDF neurons, trained on the HH *V*_1_(*t*), are in the neuron segment (red). *N_c_* = 500 for the DDF neurons.

This network segment result indicates that, indeed, we may replace the more complex HH neuron voltage activity with the reduced dimension, biophysically trained, V(t) DDF neuron in synaptic connections occurring in a network of neurons.

## 10 Summary and Discussion

This paper is a melding of many ideas from nonlinear dynamics and applied mathematics to the goal of constructing biophysically based models of observables in neurobiology. These data driven models encode the full information in experimental observations on the complex system, the neuron, which is observed in a current clamp experiment, i.e. one with a given *I_stim_*(*t*) with membrane voltage V(t) observed, and permit forecasting/predicting the voltage response of the observed neuron to other stimuli.

The method is called Data Driven Forecasting (DDF) and the model construction produces a discrete time, nonlinear map *V*(*t*) → *V*(*t* + Δ*t*) = *V*(*t* + *h*) that may be used in a network with synaptic or gap junction connections as the dynamical function of each is determined by the voltages of the presynaptic and the postsynaptic cells. We have shown this in a simple network for gap junction neuronal connections and for a network segment where a presynaptic neuron, stimulated by *I_stim_*(*t*), drives a postynaptic neuron via an excitatory synapse. The formulation of the required dynamical map for the gating variables of synaptic connections is given in Appendix B.

The DDF based network permits a computationally efficient network model where the trained DDF discrete time map, trained on biophysical data, replaces complex Hodgkin-Huxley models typically used in contemporary research. The computational advantage is achieved in two ways: (a) no differential equations at the network nodes must be solved, and (b) the nodal models are substantially reduced in complexity as it is only the observable, V(t) that is forecast.

In the timing comparisons we performed on the use of a HH NaKL model neuron solved by a standard fourth order Runge-Kutta ordinary differential equation solver compared to the forecasting efficiency of the V(t) alone DDF for the same time period with the same *I_stim_*(*t*) we found a computational improvement by a factor of 3.7. As the HH NaKL model evaluates four state variables {*V*(*t*), *m*(*t*), *h*(*t*), *n*(*t*)} while the V(t) trained DDF neuron gives only the time course of V(t): *V*(*t*) → *V*(*t* + *h*), a factor of about four in execution of the DDF is sensible.

For the data in Fig. (3) a detailed Hodgkin-Huxley model with 14 state variables, V(t) and 13 gating variables was constructed using methods of data assimilation [27]. The improvement in computational efficiency in forecasting V(t) using the DDF neuron trained on these data over the HH model was approximately 10. This suggests the idea that for a HH neuron with *N_gating_* variables, the computational efficiency of a DDF neuron accurately forecasting the membrane voltage will be about *N_gating_* faster than solving the HH model at each node of a functional network.

One must recognize that while much is gained by the DDF neuron construction, something is set aside, and that is the knowledge of the biophysics in detailed HH models of individual neurons including the operation of gating variables for the ion channels selected for the model, parameters determining the dynamics and strength of those ion channels, and other unobserved, yet relevant biophysical properties of the neurons.

In the present paper we have presented the formulation of the DDF method. Many familiar with its ingredients will recognize the provenance of such a strategy. We have also shown how this approach can be applied to experimental data from current clamp experiments from an avian songbird preparation.

The challenges ahead in building large realistic functional network models in neurobiology are significant. Again, as the DDF models are based on observations of V(t) from experimental data, the biophysical content of each representation of neuron dynamics is equivalent.

While we certainly have not shown this in the present paper, it is not inappropriate to state that the connectivity of a realistic functional network with DDF neuron models at the nodes can be established using methods of data assimilation [38, 21, 27, 2].

Data assimilation for such a network requires a model of the connectivity among the component neurons along with data on the operation of many neurons in the intact network. The ability to use Ca fluorescence experimental results [33, 23, 39], where a large number of neurons are observed, in a data assimilation environment suggests a path forward for this kind of network connectivity construction.

Another potential use of DDF arises when one does have detailed HH model neurons for network components. This may be of value in answering questions more intricate than computationally efficient neurons at the nodes of large networks. [9] and ([2], Chapter 9). It is likely that data assimilation will have been used to create such detailed biophysical models. Using V(t) from such models one can construct a reduced dimension DDF *V*(*t*) → *V*(*t* + *h*) realization of the biophysics in the HH model, and then, as we showed, one may replace the complex HH model with the reduced dimension DDF construct. One can expect to achieve computational efficiency and allow the exploration of larger biological networks when using the DDF construction to capture the neuron biophysics.

## Acknowledgements

We acknowledge support from the Neurological Disorders and Stroke Institute, National Institutes of Health, “From ion channels to graph theory in sensorimotor learning,” Grant U01 NS115821-01, and the Office of Naval Research, Grants N00014-20-1-2580 and N00014-19-1-2522. Discussions with Erik Bollt and Daniel Gauthier were instrumental in initiating this work. C. Dan Meliza (University of Virginia) and Daniel Margoliash (University of Chicago) kindly provided us with the current clamp data from experiments on neurons within the nucleus HVC of zebra finch songbirds. Discussions with Stefan Luther and Ulrich Parlitz gave us insights on how DDF may have broad applications in biophysics. Two anonymous referees assisted in improving earlier drafts of this paper.

## A Appendix A

### A.1 Guide to the Use of Radial Basis Functions in Data Driven Forecasting

To assist the reader to have an overview of how we construct and use the representations of the vector field for the DDF discrete time maps constructed from observed data we present in this Appendix a ‘manual’. Our manual is more general than the applications in the main body of the paper which has as its focus the use of the methods in neuroscience. In this ‘manual’ We will try to identify what is special for neuroscience and what is useful in general.

1. Collect D-dimensional data for the observations on your system of interest. We call these data **u**(*t_n_*) = **u**(*n*)∈ *R^D^*; *t_n_* = *t*_0_ + (*n* – 1)*h*, *n* = 1, 2, … *N*. We wish to use these data to construct a discrete time map **u**(*n*) → **u**(*n* + 1) which we then use for *n* > *N*, namely to forecast in the dynamical development of the observations beyond the data that has been collected.
2. Select a training subset of the data {**u**(*n*)}; *n* =1, 2, …, *N_T_*; *N_T_* < *N*.
3. Select a subset of the observed data to use as ‘centers’ **u**^*c*^(*q*) *q* = 1, 2, …, *N_c_*; *N_c_* ≤ *N_T_*. The RBF was designed to be an accurate interpolating function in **u** space, and the centers tell us which samples of the distribution in **u** space we utilize.
4. Select a radial basis function (RBF) ***ψ***([**u** – **u**^*c*^(*q*)]^2^, *σ*). All of RBF’s have a ‘shape factor’ *σ* that is associated with the distance in D-dimensional **u** space over which the influence of the observed data is important [40]
5. Represent the vector field **f**(**u**(*n*), **χ**) for the discrete time dynamics **u**(*n* + 1) = **u**(*n*) + **f** (**u**(*n*), **χ**). The **χ** are a set of parameters which we will learn while training the RBF.
6. The representation of **f** (**u**, **χ**) is

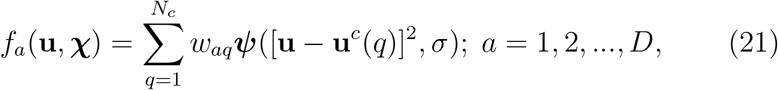

and among the parameters in **χ** are {*w_aq_*, *σ N_c_*} and others we identify in a moment. A more general RBF formulation [22, 32] allows for a polynomial in **u**, but we have not used this freedom in this paper.
7. Train the elements of **χ** via

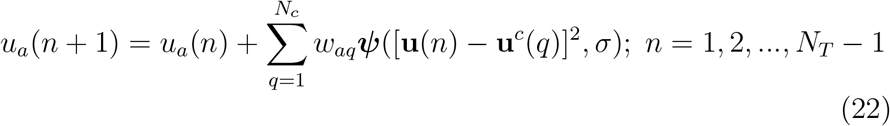
8. This training involves the inversion of a rectangular *N_c_* × *N_T_* matrix ***ψ***([**u**(*n*) – **u**^*c*^(*q*)]^2^, *σ*) which is an ill-posed problem and requires regularization [29] to make its inverse well defined. This procedure is also called *ridge regression* in the literature. The regularization consists of realizing the determination of the {*w_aq_*} as the minimization of

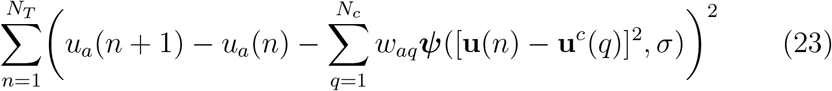

to which we add a regularization term 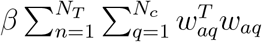. Writing **Z**_*nq*_ = ***ψ***([**u**(*n*) – **u**^*c*^(*q*)]^2^, *σ*) and *y_a_*(*n*) = *u_a_*(*n* + 1) – *u_a_*(*n*) the solution to the minimization of Eq. (23) is written as

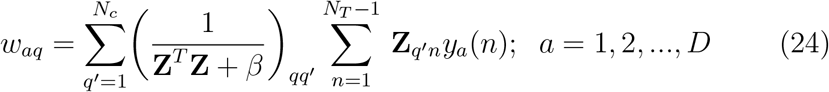 This enlarges the list of quantities in we are required to estimate to {*w_aq_*, *σ N_c_*, *β* }.
9. If one is using the time delay embedding space to ‘unproject’ the D-dimensional measurements into a larger space where the dynamics operates, then we need to estimate two more parameters: the time delay *τ* and the dimension *D_E_* of vectors **S**(*n*) in the larger space. For *D* = 1, the case we encountered in application of DDF to neuroscience problems, where *u*(*n*) = *V*(*n*), the time delay vectors are

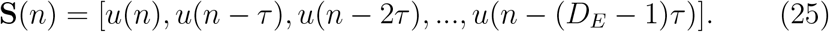 If *D* > 1, the time delay vectors will have components built with time delays of the observed **u**(*n*). The full set of parameters we need to estimate in this practical setting is now {*w_aq_*, *σ*, *N_c_*, *β*, *τ*, *D_E_*}.

To implement this protocol, we have proceeded in the following way:

There are many RBF’s to choose from. See Table 1 in [32] for a list. We have not tried them all. We have primarily used the Gaussian

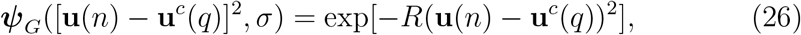

in our calculations. R is the precision of the Gaussian.

We also used the multiquadric of Hardy [13, 26, 22, 32]

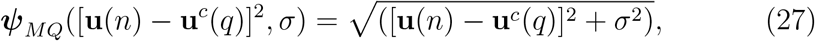

and found on some problems essentially equivalent results. In what follows, and in the work in this paper, we remain with ***ψ***_*G*_([**u**(*n*) – **u**^*c*^(*q*)]^2^, *σ*).

### A.2 Including Polynomial Terms in the Representation of f (**u**)

It can be helpful is to include polynomial terms as well as RBF’s in the representation of the discrete vector field [32][22]. The vector field representation becomes:

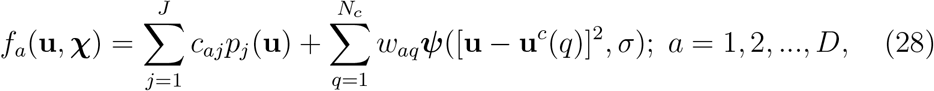

which is what we see in Eq. (1).

The addition of polynomials is indicated if we know on other grounds something about the underlying dynamical of the source of the data that suggests polynomial terms in the vector fields are present. That is not the case in applying these methods to neuroscience, so except for the term **u**(*n*) in Eq (22), we do not use the freedom of further polynomials in **u**.

### A.3 How to choose Centers

The choice of centers, the **u**^*c*^ vectors in the RBF, is constrained by the data points in the training window 1 ≤ *n* ≤ *N_T_*. We used K-means clustering [10] to select the **u**^*c*^, and this is a way of selecting centers where they are needed to sample the distribution in **u** space. While there may be more advanced methods, we have found K-means to work well enough and have used the method in finding all of the results in the paper.

Another aspect to consider is the number of centers *N_c_*. A general rule of thumb is that more centers can better, as the distribution in **u** space is better sampled. However, this comes at the cost of both memory and computational time. In this paper, we typically chose about 5000 centers for training lengths *N_T_* ≈ 25 – 50 × 10^4^.

There is a possible strategy with regard the selecting *N_c_* that balances a fine grained sampling of the distribution in **u** space against the increased computing cost in evaluating the dynamical map **u**(*n*) → **u**(*n* +1) when *N_c_*. When presented with a data set that is noisy the resolution in **u** space is coarse grained when the data arrives. The idea is to recognize that limitation on how well any DDF can perform by evaluating a metric on the match between the predictions of the trained DDF {**u**_*DDF*_(*n*) compared to the known, noisy knowledge we have of the data **u**_*data*_(*n*) for *n* ≥ *N_T_*. This could be something like 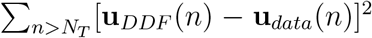. Noise will limit the improvements in forecasting quality as *N_c_* increases, and the performance of the metric should lead to a modest *N_c_* larger than which no increase in forecasting skill is seen.

#### A.3.1 Centers and RBF’s

We found that the obvious choices for DDF worked out well, namely choosing Gaussian RBF’s and K-means clustering. These strategies are commonly used in the RBF literature.

However, these are far from the only good working ideas there and others could possibly work better. The centers could also be chosen by emphasizing places where the first or second derivatives in the data **u**(*n*) are greatest. This could work well because it could select more centers where data might be sparse, for example when the neuron data is spiking and regions are traversed at higher speed than in sub-threshold regions with smaller derivatives in **u**(*n*) [30].

### A.4 Working with Imposed Driving Forces Acting on the Source of the Data {u(*n*)}: Stimulating Currents *I_stim_*(*t*)

We have not found a general formulation for DDF in the case of a driven dynamical system. One natural idea is to include those forces, call them **K***_ext_*(*t*) on the list of observed quantities in the data: {**u**(*n*)} → {**u**(*n*), **K**_*ext*_(*n*)} and enter this into the arguments of the relevant RBFs.

In many interesting situations in Physics and Neurobiology, however, the forces are additive, so the formulation presented in this paper maybe applicable to many interesting problems.

If we were analyzing the underlying detailed dynamical equations of the processes producing the data, the *K_ext_*(*t*) would have to be provided as they are not given a differential equation of their own, but specified by the user. This is true when we solve the differential equations for a model of the driven systems and remains true for DDF.

In the case of the dynamics of neurons, we are in a fortunate situation. In this instance the forcing is from currents external to the intrinsic currents associated with the ion channel dynamics. The voltage HH equation is a statement of current conservation and currents are additive.

This permitted us to write Eq. (4)

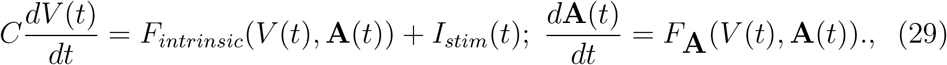

and is an example where some knowledge of the biophysics or other Physics of the data source is of value.

### A.5 Evaluating the χ

We have two classes of parameters in our representation of the vector field **f**(**u**, **χ**): those coefficients {*c_aj_*, *w_aj_*} which weight the polynomial and RBF contributions in Eq. (1). These are determined by linear algebra from

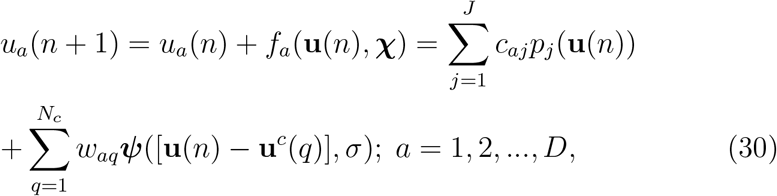

when the parameters **Ξ** = {*R*, *β*, *τ*, *D_E_*} are fixed.

We proceeded by selecting a set of **Ξ**, making a coarse grid search over the **Ξ**, performing the regularized linear algebra to train the coefficients ***φ*** = {*c_aj_, w_aj_*} for each choice of the **Ξ**, and using this trained representation to perform a validation test for each choice. **χ** = {***φ*, *ξ***}.

To accomplish this validation step this we selected the unused part of the known data with *N_V_* = *N* – *N_T_* elements and calculated a validation set of forecast values **u**^*V*^ (*n*)

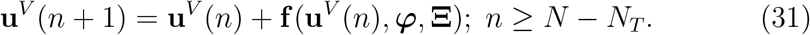

The we compared the values **u**^*V*^(*n*), which are dependent on the choice of **Ξ**, with the known data. Effectively we evaluated the cost function

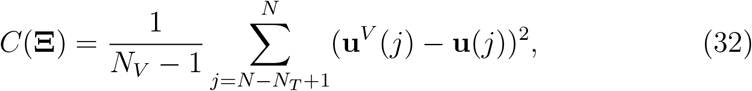

and sought a minimum over the **Ξ**.

Our practice has been to examine the quality of **u**^*V*^(*n*) ≈ **u**(*n*), locate regions of the **Ξ** where they match well, and refine the grid search in the **Ξ** as required.

#### A.5.1 Differential Evolution to search for the minimum of *C*(Ξ)

Differential Evolution is a genetic algorithm that could be implemented in our DDF construction to improve our ability to create DDF models by more precisely choosing sets of parameters **Ξ** = {*R, β, τ*, *D_E_*}. The method is described in [35].

It works by initiating a parent set of parameters from a user defined uniform distribution. From that start, new “children” sets of **Ξ** are made from combining the parents in a algorithmic way and comparing the new validation forecast {**u**^*V*^(*j*)} to the old one. A user defined cost function, such as Eq. (32), is used to compare the the children to their parents.

As generations go by, parents will gradually be replaced by children until all the parents converge on a minimum in the cost function. If the user is able to define a clever cost function, they could get a result that hope to surpass those from the simple grid searching employed so far

### A.6 Code for Our Implementation of DDF

We did all of our testing and programming in Python. We give links to the code we used published in GitHub. In the links we include DDF code that was written purely for Radial Basis Functions, in the repository is code for both Gaussian and MultiQuadric forms of the RBF. There is a link to a DDF neuron repository that includes the code that has been built specifically for the study of an NaKL neuron where only the voltage and the stimulus is known; this means that the code is built to perform time delay embedding on a single dimension of voltage. The treatment of the stimulus current in this code is as was described earlier. The third and final attached link is to a method of using DDF with polynomial terms in the representation of **f** (**u**, **χ**) as an additional resource to the reader.

- https://github.com/RandarserousRex/Data-Driven-Forecasting-Radial-Basis-Function-Method
- https://github.com/RandarserousRex/DDF-Applications-to-Neurons
- https://github.com/RandarserousRex/Data-Driven-Forecasting-Taylor-Method

### A.7 Memory Management

An important note to consider is the size of the matrices involved in training, for they can grow to the size of gigabytes. For example, the largest matrix involved in training is that of all values of the RBF’s at all times, *N_T_* by *N_c_*, can have a standard length of 25,000 data points with 5,000 centers. Typically we use float64 for all our values resulting in this matrix having a size 25, 000(*N_T_*) × 5, 000(*N_c_*) × 8 = 1 Gigabyte. We need another Gigabyte for its transpose used in the regularized matrix inversion calculation. This could be a limiting factor if one is running multiple tests in parallel on a CPU or on a cluster with limited storage.

### A.8 Parallelizing DDF Calculations

This section serves as a suggestion to take advantage of the ability to run DDF code in parallel. The grid searching method discussed above lends itself readily to parallel programming. Even the Differential Evolution method can be run in parallel. A single trial of DDF doesn’t have much potential for parallel operations as the training is just matrix multiplication, and the forecasting is a step by step process that relies on the result of the previous step.

## B Appendix B

In the main text we implemented a simple two neuron circuit with gap junction connections between the neurons. A richer network consists of more neurons and allows for synaptic connections as well. Starting with an equation for synaptic currents *I_synaptic_*(*t*) we need a discrete time map to update the time dependence in a map for V(t). This is given now.

### B.1 Discrete Time Synaptic Dynamics

In the discrete time dynamical rules for the membrane voltages of the neurons in a network we require the gating variable at discrete time *t_n_* and *t_n_* + *h* = *t*_*n*+1_.

The synaptic current between a presynaptic neuron with voltage *V_pre_*(*t*) and a postsynaptic neuron with voltage *V_post_*(*t*) is described by

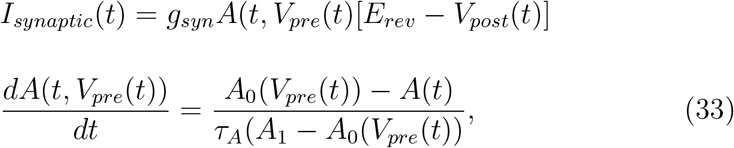

where *A*(*t*, *V_pre_*(*t*)) is a synaptic gating variable. It is opened *A*(*t*, *V_pre_*(*t*)) ≈ 1 when neurotransmitter binds onto receptors on the postsynaptic cell. It is closed *A*(*t*, *V_pre_*(*t*)) ≈ 0 when that neurotransmitter is released from the postsynaptic receptor.

Henceforth we denote the gating variable *A*(*t*, *V_pre_*(*t*)) as *A*(*t*) for brevity.

In Eq. (33) we have time constants *τ*(*A*_1_ – *A*_0_(*V_pre_*(*t*)) and a function *A*_0_(*V_pre_*(*t*)) to specify. The function *A*_0_ (*V*) described the transition of the gating variable from *A*(*t*) ≈ 0, namely a closed synaptic channel, to *A*(*t*) ≈ 1, an open synaptic channel.

We represent this transition in the neighborhood of a transition at a voltage *V*_0_ from closed to open by writing

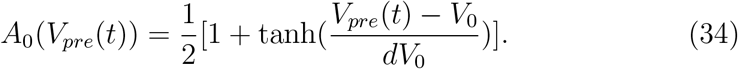

This function moves from very near 0 when *V* ≪ *V*_0_ to very near 1 when *V_pre_*(*t*) ≫ *V*_0_, as desired, and it does so over an interval in voltage *dV*_0_.

When the width of this transition *dV*_0_ is small, *A*_0_(*V*) is essentially a step function in voltage.

When *V_pre_*(*t*) < *V*_0_ Eq. (33) is approximately

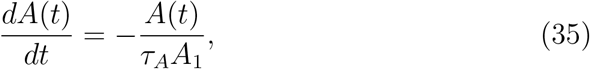

and when *V_pre_*(*t*) > *V*_0_ Eq. (33) is approximately

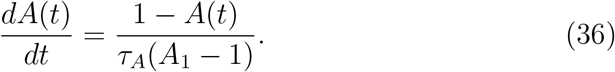

By integrating each of these equations from time t to time t+h, we find for *V* ≪ *V*_0_

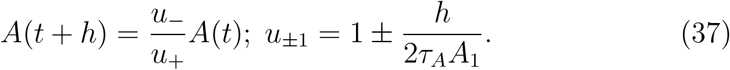

While for *V_pre_*(*t*) ≫ *V*_0_ we find

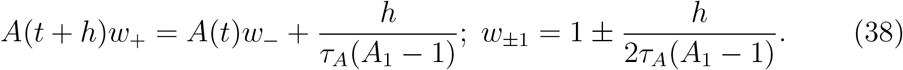

To avoid the discontinuous change in voltage associated with a step function in *V_pre_* we write a smoother transition over an interval *dV* for the synaptic gating variable *A*(*t*):

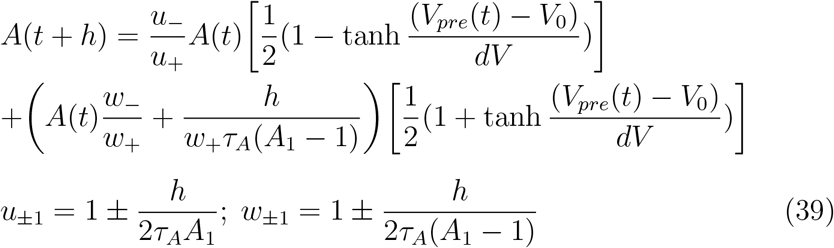

### B.2 Excitatory and Inhibitory Synaptic Formulation

In Eq. (33) we have two time constants {*τ_A_A*_1_, *τ_A_*(*A*_1_ – 1)} corresponding to the time it takes for a neurotransmitter to undock from a postsynaptic receptor and to dock at the receptor, respectively.

For an *excitatory* AMPA receptor, mediated by the neurotransmitter glutamate, the approximate value of *τ_A_A*_1_ ≈ 3ms, and of *τ_A_*(*A*_1_ – 1) ≈ 1ms. So for this excitatory synaptic connection *τ_A_* = 2 ms and *A*_1_ = 1.5. For an *inhibitory* synaptic connection, mediated by GABA, *τ_A_A*_1_ ≈ 8 ms, and of *τ_A_*(*A*_1_ – 1) ≈ 2ms, leading to *τ_A_* = 6 ms and *A*_1_ = 4/3.

## C Appendix C

### What waveforms should be chosen for *I_stim_*(*t*)?

This question arose in the valuable comments of the reviewers and is certainly an important issue in training a DDF model neuron so that it will respond accurately to any stimulating current it may encounter in a network or from environmental stimulation.

The fact that the stimulating current enters the biophysical (HH) equations in an additive manner, reflecting the way current conservation works in all electric circuits, suggests that a judicious choice of *I_stim_*(*t*) is possible.

We do not have a mathematical statement about what stimulating currents should be used to train a DDF, or for that matter a HH neuron model via data assimilation. However, we do have a biophysical argument.

1. If one wanted to learn how a DDF would respond when parameter value is changed, then presenting data to the DDF training protocol from a range of parameter values would permit the DDF to interpolate through that range. DDFs, and radial basis functions (RBFs) are interpolating mathematical devices by construction. With enough data sampling enough of the dynamical system attractor the RBF can accurately interpolate among the given data points.
2. As we have no differential equation for the stimulating current *I_stim_*(*t_k_*)); *t_k_* = *t*_0_ + *hk*, k = 0, 1, 2, … N are just like parameters in the voltage or any other DDF. If we choose a training I(t) which evokes a V(t) over the dynamical range expected of a neuron, say −100 mV to 100 mV, then we will be able to interpolate to new V(t) values within the training range. That is likely not to be quite enough as *I_stim_*(*t*) is also a time series, albeit prescribed by the user, so we have found that the strength of *I_stim_*(*t*) in its Fourier power spectrum must lie in a range which can be determined by the data it is presented. Biophysically, if the *I_stim_*(*t*) frequencies are too high, they would be filtered out by the cell membranes RC time constant, as that acts as a low pass filter.

Just this line of thinking led to the stimulating currents one sees in each of the experimental time courses presented in the paper. Before we used this biophysical heuristic, data assimilation—training a HH model with data—was unsuccessful. With this heuristic guiding the choices of *I_stim_*(*t*), successful DA is now routine.

## Notes

https://github.com/RandarserousRex?tab=repositories

### Competing Interest Statement

The authors have declared no competing interest.

### Summary of Updates

An Additional Section 9.2 was added to include the test of a two neuron network of ligand gated synaptic connections. An Appendix Section C has been added to discuss what stimulus are appropriate. General clean up has been performed as well

